# Multi-stage-mixing to control the supramolecular structure of lipid nanoparticles, thereby creating a core-then-shell arrangement that improves performance by orders of magnitude

**DOI:** 10.1101/2024.11.12.623321

**Authors:** Jia Nong, Xijing Gong, Quang Minh Dang, Sachchidanand Tiwari, Manthan Patel, Jichuan Wu, Andrew Hanna, Wook-Jin Park, Elena N. Atochina-Vasserman, Hui-Ting Huang, Oscar A. Marcos-Contreras, Kahlilia C. Morris-Blanco, Jonathan J. Miner, Drew Weissman, Vladimir R. Muzykantov, Kushol Gupta, David Issadore, Jacob W. Myerson, Zhicheng Wang, Jacob S. Brenner

## Abstract

As they became the dominant gene therapy platform, lipid nanoparticles (LNPs) experienced nearly all their innovation in varying the structure of individual molecules in LNPs. This ignored control of the spatial arrangement of molecules, which is suboptimal because supramolecular structure determines function in biology. To control LNPs’ supramolecular structure, we introduce multi-stage-mixing (MSM) to successively add different molecules to LNPs. We first utilize MSM to create a core-then-shell (CTS) synthesis. CTS-LNPs display a clear core-shell structure, vastly lower frequency of LNPs containing no detectable mRNA, and improved mRNA-LNP expression. With DNA-loaded LNPs, which for decades lagged behind mRNA-LNPs due to low expression, CTS improved DNA-LNPs’ protein expression by 2-3 orders of magnitude, bringing it within range of mRNA-LNPs. These results show that supramolecular arrangement is critical to LNP performance and can be controlled by mixing methodology. Further, MSM/CTS have finally made DNA-LNPs into a practical platform for long-term gene expression.

## INTRODUCTION

The billion-patient success of the COVID-19 mRNA lipid nanoparticle (LNP) vaccine has led to >>$10B investment by industry and governmental funders in developing LNPs as vaccines and therapeutics [1]. This has accelerated the innovation of LNPs, which had already been iteratively improving for 20+ years. Nearly all that innovation has gone into varying the structure of individual lipid molecules in LNPs, especially LNPs’ ionizable lipids [2, 3].

This laser focus on individual molecules’ structures has largely ignored the role of “supramolecular structure”, meaning the spatial arrangement of molecules within LNPs. Supramolecular structure is likely important to the future of LNPs, as nearly all biological nanoparticles have highly ordered arrangement of their molecules. Examples include humans’ endogenous nanoparticles like high density lipoprotein (HDL), which forms highly ordered discs; similarly, viruses display complex arrangements of diverse molecular components, with very low variability in supramolecular structure between individual virions. In stark contrast to viruses’ highly ordered supramolecular structures are current LNPs, which are amorphous / glass-like and display great variability between individual particles in a given batch. Given that viruses are much more efficient at transfection than LNPs, it is worth testing if improved control over supramolecular order might benefit LNPs’ transfection efficiency. Further, we previously showed that the side effects of therapeutic nanoparticles can be very dependent on supramolecular structure; for a particular class of nanoparticles (those with surface proteins), amorphous nanoparticles strongly activated the “complement cascade” (and its downstream side effects), while more ordered, near-crystalline nanoparticles evaded complement [4]. Thus, there are multiple compelling reasons to control the supramolecular structure of LNPs, ranging from transfection efficiency to side effects.

Indirect control of LNPs’ supramolecular structure has been achieved to some extent in a couple cases by varying the chemical species that go into LNP synthesis; namely, inclusion of unusual neutral lipids and high concentrations of small ions. For example, glycerol monooleate (GMO) can promote particular lyotropic liquid crystalline phases in LNPs and improve transfection efficiency 2-5-fold [5]. Similarly, increasing the ion concentration in buffers during LNP synthesis can increase the frequency of “bleb” structures in LNPs, which correlates with improved transfection efficiency for low-potency ionizable lipids [6]. In both cases, the changes in supramolecular structure came from the addition of a particular chemical species, rather than methods of actively arranging a given set of molecules.

Theoretically, a direct way to control supramolecular structure, without varying the constituent molecules, is via innovations in the synthesis methods of LNPs. The synthesis of LNPs involves mixing of constituent molecules (lipids, nucleic acids, small ions, etc.), incubation steps (varying temperature, time, etc.), and purification. Unfortunately, very little innovation effort has been put into improving LNP performance metrics by redesign of LNP synthesis methods, especially the most important step, mixing.

Indeed, LNP mixing methodology has remained largely the same for decades. For the last twenty years, LNPs have been produced by rapidly mixing two liquid phases: an aqueous phase containing dissolved nucleic acids (almost always RNA); and an organic phase (usually ethanol) in which lipids are dissolved (**Supplementary** Fig. 2A). Initially, these two liquids were mixed in microfluidic devices, but later moved to much larger mixing chambers, such as confined impinging jet (CIJ) mixers, which aided with industrial scale-up manufacturing with the COVID vaccines [7]. The fact that LNPs made via microfluidics (using laminar flow) were indistinguishable from macro- scale CIJ mixers (employing turbulent mixing) was surprising, and suggested that the method of mixing LNPs does not affect LNP performance. This idea that mixing methodology is irrelevant to LNP characteristics was further supported by an influential paper that showed that LNP performance was also barely affected by changing to “low-tech” methods of mixing such as vortexing and even hand-pipetting [8]. Thus, the mixing method has largely been ignored as a route of improving LNP performance.

Here we revisited the idea of innovating the mixing methodology of LNPs, and found that such innovation could improve LNP performance by orders of magnitude. We hypothesized that we could control LNP’s supramolecular arrangement of molecules by multi-stage mixing (MSM), in which different molecular species are sequentially added to a growing LNP structure. We initially tested whether MSM could make the simplest possible pre-specified supramolecular structure, two layers. To enable this, we devised microfluidics that could perform a core-then-shell (CTS) synthesis: first forming a “core” comprised of one set of molecular species, and then a “shell” composed of other molecular species. In a CTS synthesis of LNPs, there are two stages of mixing (**Fig. 1A**). First, nucleic acid (anionic) is mixed with a cationic molecule, and these two species electrostatically condense to form a “core” (**Fig. 1A**). Microseconds later, the cores are mixed with a third organic solvent stream containing the standard LNP lipids (ionizable lipid, cholesterol, “helper” phospholipid, PEGylated lipid), forming a “shell” around the cores. Importantly, without the shell, these cores are metastable, aggregating to >2000 nanometers within minutes, rendering free “cores” unfeasible for *in vivo* use. MSM captures such metastable intermediates, which would be impossible to achieve through equilibrium processes. This capture of metastable states points out a fundamental difference between MSM and traditional layer-by-layer (LbL) techniques. Traditional LbL involves sequential deposition of oppositely charged materials onto a preformed, thermodynamically stable core [9]. Further, LbL requires multiple separation and purification steps between layer additions, which lead to material loss, batch-to-batch variability, and no industrial scale-up after decades. By contrast, MSM enables capture of favorable metastable states, while maintaining continuous flow production, with enhanced reproducibility and scalability.

**Figure 1:**
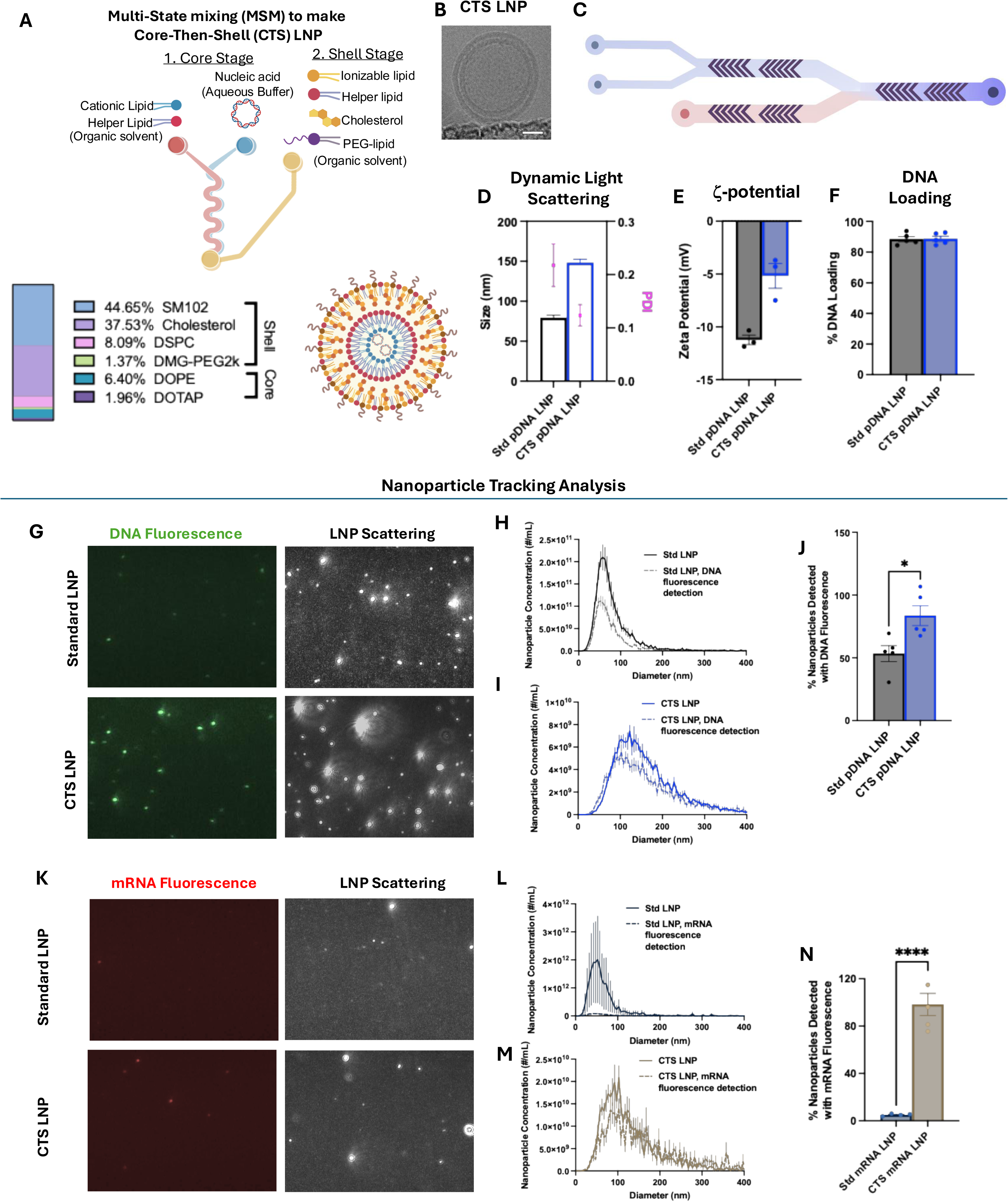
Multi-stage mixing (MSM) to produce core-then-shell (CTS) improves the encapsulation of nucleic acids. A) Schematic illustration of making CTS LNP. Instead of the standard single mixing stage LNPs have used for decades, CTS LNPs are made via multi-stage mixing. First, in the Core Stage, the DNA (aqueous buffer) is mixed via a microfluidic with an organic solvent containing a cationic lipid and helper lipid. This condenses the DNA into a “core.” Immediately upon core formation, the cores are mixed with standard LNP lipids in the organic phase, to form the “shell”. **B) Representative Cryo-EM** shows that CTS-LNPs created using multi-stage mixing (MSM) display distinct core and shell phases. The white bar shown denotes a length of 25 nm. **C) Schematic illustration of in-house made microfluidic chip. D) Dynamic Light Scattering (DLS).** CTS LNPs are larger than standard (Std) LNPs, but have a lower polydispersity index (PDI). **E) ζ-Potential Analysis using ZetaSizer.** CTS LNPs are closer to neutral, measuring the bulk nanoparticle population. **F) DNA entrapment efficiency:** CTS LNP and standard LNP have similar DNA entrapment efficiency around 90%. **G, H, I) DNA cargo payload distribution in LNPs.** To evaluate the distribution of DNA within a population of LNPs, we used Nanoparticle Tracking Analysis (NTA) to trace particles using light scattering (right panels) and simultaneously measure DNA loading into each particle via fluorescently labeled DNA (left panels). In H & I, we plot the distribution of nanoparticle sizes, as measured by light scattering (solid lines) or DNA fluorescence (dashed). H shows that in standard LNPs, a low fraction of particles contain detectable DNA, while I shows that for CTS LNPs a very high fraction of nanoparticles contain DNA. **J) quantification of H-I**, showing that in standard LNPs, only ∼50% of LNPs have detectable DNA, while 83% of CTS LNPs have detectable DNA. **K, L, M) RNA cargo payload distribution in LNPs.** NTA evaluation of mRNA payload in standard LNP vs CTS LNP shows similar trend as in DNA LNP. **N) Quantification of L-M**, showing that in standard LNPs, only 5% of LNPs have detectable RNA, while 98% of CTS LNPs have detectable RNA.

Here we begin by showing that MSM and CTS do indeed produce LNPs with radically different supramolecular structure than standard LNPs, including confirming that the CTS synthesis technique produces the hypothesized core-then-shell structure. Further, we showed that the mixing methodology, not just the lipid formula (molar ratios), is essential to the supramolecular structure, as switching between CTS vs standard two-stream synthesis produces particles with extremely different distributions of surface charge.

Next we showed that CTS produces LNPs with many advantages. First, we showed that CTS decreases the fraction of LNPs in a batch that are “empty” (have undetectably low nucleic acid levels), which is important since we had previously shown that “empty” LNPs are more inflammatory [10]. Second, we showed that CTS LNPs loaded with mRNA express proteins at levels up to 4x higher standard LNPs. However, the benefits of MSM and CTS are best shown with DNA-loaded LNPs (DNA-LNPs).

DNA-LNPs have long been a top goal in genetic medicine, as they could solve the two fundamental limitations of mRNA-LNPs: i) mRNA’s half-life is just hours, which precludes treatment of chronic diseases [11]; ii) mRNA has no promoter region, so an inability to control cell-type-specific protein expression, shifting most LNP-stimulated protein expression to off-target cells [12]. DNA can solve both problems, with its yearslong intracellular stability and cell-type-specific promoters. Unfortunately, in 20 years of LNP development, DNA-LNPs never advanced because they suffered two problems: a) DNA-LNPs are toxic, killing mice within 2 days; b) DNA-LNPs’ protein expression is orders of magnitude less than that of mRNA-LNPs. We solved the first problem in a paper recently accepted at *Nature Biotechnology* [13]. We showed that the toxicity of DNA-LNPs is caused by detection of cytosolic DNA by cGAS-STING. Next, we loaded DNA-LNPs with nitrated lipids, which are mammals’ natural negative feedback inhibitor of STING. DNA-LNPs loaded with such STING-inhibiting lipids eliminated DNA-LNP mortality. Further, these STING-inhibiting DNA-LNPs stably express protein *in vivo* for at least 3 months per dose.

Here we use MSM & CTS to solve DNA-LNPs’ second problem, that their protein expression levels are 1,000- to 10,000-fold lower than the peak of mRNA-LNPs. We hypothesized that CTS could pack DNA into tight “cores” that would protect the DNA from DNases and cytosolic DNA sensors, and enable easier transit of DNA through the narrow pores of the cytosolic gel and the nuclear pore complex [14–16]. We show that CTS does, indeed, improve DNA-LNP expression by 2-3 orders of magnitude, allowing DNA-LNPs to drive protein expression similar to that of mRNA-LNPs at their peak. Thus, MSM & CTS have finally made DNA-LNPs into a practical tool for genetic medicine. In the Discussion, we explain how DNA-LNPs can now bring genetic medicine to common chronic diseases, and this was all made possible because MSM allows control of LNPs’ supramolecular structure.

## RESULTS

### Multi-stage mixing (MSM) / core-then-shell (CTS) synthesis allows control over LNPs’ supramolecular structure, and improves the uniformity of nucleic acid loading

To control the supramolecular structure (spatial arrangement of molecules) of LNPs using MSM, in this study we start with the simplest possible arrangement of molecules within an LNP: two layers, with each layer composed of different molecular species. As LNPs are spheres, a two-layer arrangement equates to a “core” and a “shell.” Thus, the simplest MSM protocol is a synthesis that forms a core and then a shell (“core-then-shell” synthesis [CTS]). In CTS synthesis, in Mixing Stage 1, nucleic acid (in aqueous solution) is mixed with a cationic species, leading to electrostatic condensation to form a “core” (**Fig. 1A**). In our first example of CTS, the cationic species is the lipid DOTAP, and we additionally add a common LNP component, the zwitterionic lipid DOPE. We have subsequently accomplished CTS synthesis with many different cationic species, including ionizable cationic lipids like those typical of LNPs. Notably, these cores are not thermodynamically stable - they rapidly grow to sizes exceeding 2000 nm, making them unsuitable for both LNP incorporation and *in vivo* administration. To overcome this, in Mixing Stage 2, our CTS synthesis encapsulates these nascent cores with standard LNP lipids before they can expand, forming a stabilizing "shell" that arrests core growth. Based on lipoplex studies showing that an N:P of DOTAP:DNA ratios above 2 impede nuclear delivery of DNA, we selected a DOTAP:DNA N:P of 0.63 [17]. Notably, this leaves DOTAP at a final molar % of total lipids of only 1.96%. This is far below the 10-50% range used for DOTAP LNPs with lung tropism, which we previously showed induced thrombosis [18, 19]. Indeed, we show in **Supplementary** Fig. 1 that this low level of DOTAP, buried in the core, does not induce accidental lung tropism or clotting.

Cryo-electron microscopy confirmed successful formulation of the hypothesized core-shell architecture (**Fig. 1B**). While standard LNPs exhibited a characteristic amorphous structure (**Supplementary** Fig. 2A), CTS LNPs displayed a distinctive core-shell architecture, marked by a well-defined central region surrounded by two distinct outer shells. This shows that MSM / CTS enable control of the supramolecular structure of LNPs.

These CTS LNPs can be manufactured using either commercial dilution cartridges with 3 channels in the NanoAssemblr™ Ignite™ machine, or our in-house fabricated microfluidic devices, which are orders of magnitude cheaper than the single-use Ignite cartridges (**Fig. 1C**). For the studies presented in this manuscript, we primarily used the cartridge-based system, which produced CTS LNPs that are larger than standard DNA-LNPs but show a lower polydispersity index (PDI) (**Fig. 1D**), indicating more uniform size distribution. While the ζ-potential measured in bulk shows these CTS LNPs are less negatively charged than standard DNA-LNPs, they still maintain a slight negative charge of -5 mV (**Fig. 1E**). This is important given known toxicities of positively charged LNPs. The DNA entrapment efficiency remains high at ∼90% for both standard and CTS LNPs (**Fig. 1F**). Notably, our in-house microfluidic devices (**Supplementary** Fig. 3**, Supplementary Table 1**) produced CTS LNPs with comparable characteristics across all these parameters when screened across a range of shell lipid:DNA weight ratios, demonstrating that the multi-stage mixing (MSM) process is the key enabling factor for successful CTS LNP formation. Having established the basic physical characteristics of CTS LNPs, we next examined the uniformity of nucleic acid loading across the particle population. Recent studies have shown that within a standard LNP population, only a small fraction of the individual particles is actually loaded with detectable mRNA, while the rest are "empty" (undetectable mRNA) [20, 21]. We recently showed that empty LNPs are much more inflammatory than LNPs containing nucleic acids [10]. To assess loading uniformity at the single-particle level, we prepared LNPs with fluorescently labeled DNA and analyzed them via nanoparticle tracking analysis (NTA) on a Malvern Panalytical NanoSight. The NanoSight can trace all nanoparticles via light scattering (**Fig. 1G**, right panel), and detect whether each LNP contains detectable DNA via the fluorescent signal (**Fig. 1G**, left panel). Comparing standard LNPs (**Fig. 1H**) with CTS LNPs (**Fig. 1I**), it is evident that a higher fraction of CTS LNPs have detectable DNA in them. This is quantified in **Fig. 1J**, which shows that 83% of CTS LNPs have detectable DNA, while only 53% of standard LNPs do. We observed a similar trend in CTS LNPs loaded with mRNA (**Fig. 1K-N**). The quantification results show that more than 95% of CTS LNPs contain detectable mRNA, while only 5% of standard LNPs do. This loading fraction depends on the RNA/DNA detection threshold of the assay and is likely sensitive to lipid composition and cargo identity. However, this data shows that at least for some LNP formulations, CTS provides nucleic acid loading that is more uniform across the population of particles.

### Structural analysis reveals that core-then-shell (CTS) DNA-LNPs possess a true core and shell that is highly distinct from traditional LNPs

To directly show that the CTS protocol generates LNP structures that are distinct from standard LNPs, we characterized LNPs with single particle zeta potential measurements, differential scanning microcalorimetry (DSC), small angle X-ray scattering (SAXS), and multi angle light scattering (MALS).

We measured particle σ-potentials at the single-molecule level using an NTA device that performs microelectrophoresis. We compared two compositionally identical formulations of DNA-LNPs containing 1.96% DOTAP, prepared using different mixing methods: one with MSM microfluidics, as described in **Fig. 1A**, and the other using traditional single-step microfluidic mixing (shown in **Supplementary** Fig. 2A). The σ-potential distribution for MSM LNPs showed a broad range of moderate to large potentials like that of standard LNPs without DOTAP, ranging from 0 to -50 mV (blue and black curves in **Fig. 2A**). However, the DOTAP formulation produced with single-step mixing had two sharp peaks, one at +5mV and one at -5mV. Thus, MSM produces particles with significantly different surface charge properties compared to single-step mixing. These findings indicate that physicochemical properties, and thus supramolecular arrangment of LNPs’ molecules, are influenced not only by lipid composition but also by the mixing protocol.

**Figure 2:**
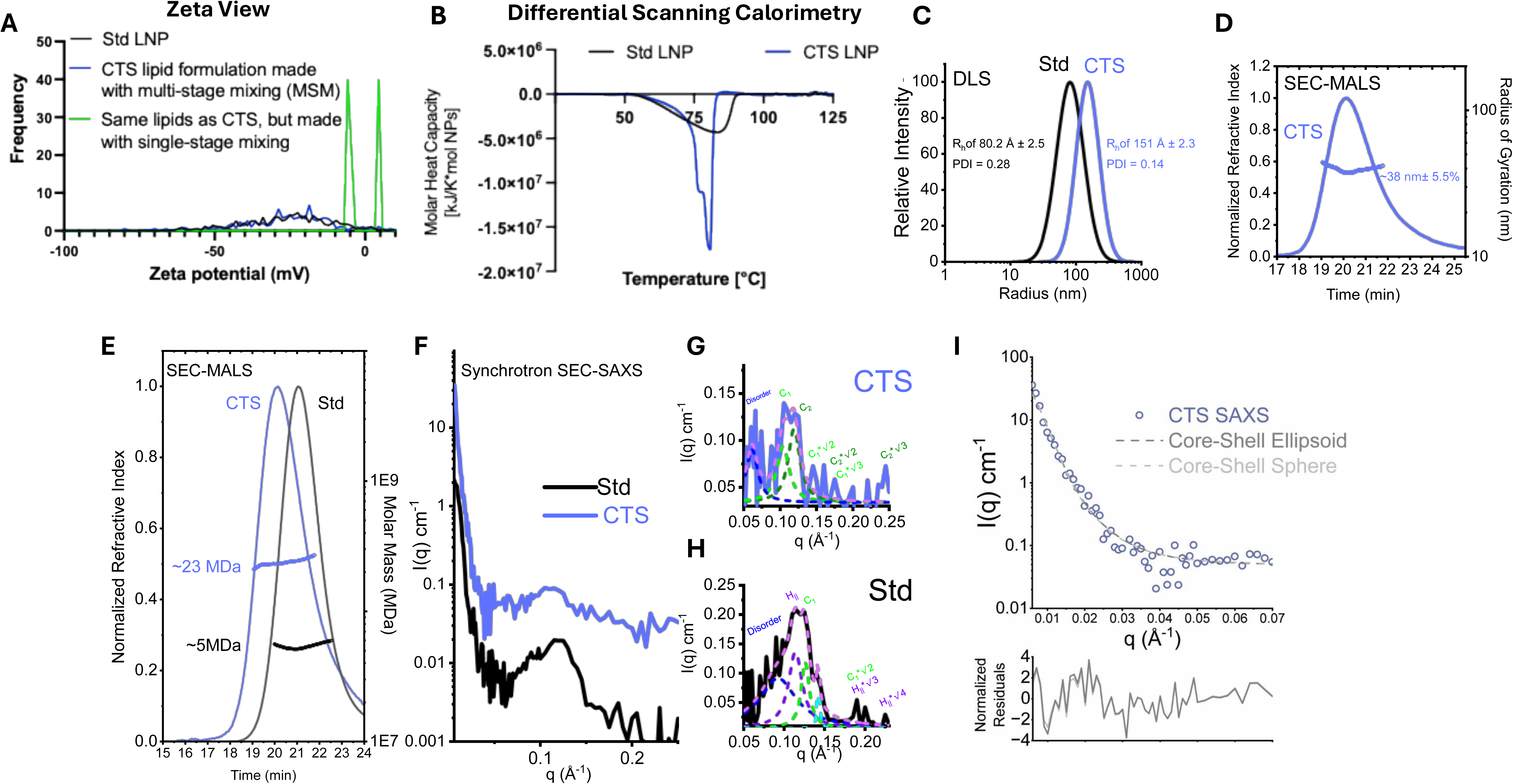
Multi-stage mixing dramatically changes the internal structure of DNA-LNPs, producing a distinct core vs shell. A) ζ-Potential Analysis. We compared the ζ-potentials of two LNP formulations: the CTS lipid formulation (containing <2% DOTAP), made either with MSM or single-stage (traditional) mixing. Single-stage mixing (green) produces two very separate populations with very low magnitude potentials, one positive and one negative; by contrast, MSM (blue) produces CTS LNPs with a single broad distribution of potentials between 0 mV and -50 mV, like seen with the standard LNP formulation (no DOTAP, black). This demonstrates that MSM is critical to creating CTS LNPs. **B) Differential Scanning Calorimetry (DSC).** The molar heat capacity of CTS and standard LNP particles were determined. Standard LNPs (black line) display a broad profile and short peak, consistent with modestly stable population. In contrast, CTS-LNPs (blue line) shows two sharp peaks with a larger magnitude, consistent with a more uniform and well-ordered population of higher stability. **C) Dynamic Light Scattering (DLS).** Particle size distributions, normalized for intensity, as determined by DLS are shown as lognormal distributions, with CTS shown in blue and standard LNPs shown in black. CTS particles had a determined hydrodynamic radius (R_h_) of 151 Å ± 2.3 (polydispersity index (PDI) = 0.14) compared to 80.2 Å ± 2.5 (PDI = 0.28) for the standard formulation. **D) Radius of Gyration Analysis (R_g_) using Size-exclusion Chromatography In-line with Multiangle Light Scattering (SEC-MALS).** Shown is a double-y plot, where the left axis provides the normalized refractive index for eluant as a function of retention time for CTS particles (blue line), while the right axis provides on a log-scale the radius of gyration (R_g_, in nm) of the CTS particles as determined using 18-angle light scattering (blue circles). Across the peak region analyzed, a R_g_ of 39.5 nm ± 5.5% was determined. A similar analysis could not be performed for standard LNP particles, as the particles were well below the size threshold needed for reliable determination of R_g_. **E) Mass profiles using SEC-MALS.** Shown is a double-y plot, where the left axis provides the normalized refractive index for eluant as a function of retention time for standard (black line) and CTS particles (blue line), while the right axis provides on a log-scale the determined molar mass. Across the peak regions analyzed, a M_w_ of 23.7 MDa ± 2.1% was determined for CTS particles and 5.4 MDa ± 3.1% for the standard LNP particles. **F) Synchrotron Size-Exclusion Chromatography in-line with Small-Angle X-ray Scattering (SEC-SAXS).** Log- linear plots of SAXS data from standard LNPs (black line) and CTS particles (blue line), showing the decay of scattering intensity as a function of the scattering vector q (where *q*=4πs θ/λ). Characteristic first-order Braggs peak features are observed between *q*=0.1-0.2 A^-1^. See Supplemental Figure 1 for additional information. **G) Multiple Lorentz Function analysis of Braggs peaks from SAXS of CTS Particles**. Shown as blue line are the SAXS data in a linear-linear plot, as a blue line. Multiple Lorentz fits were applied to deconvolute the data and are shown as dotted lines. The assignment of the cubosome phase peaks with the spacing of *d*=62.8 Å and corresponding peaks at integral values of *q*\*√2 and *√3. The overlapping peak at *q*=0.125 Å^-1^ (*d*=50.2 Å) could additionally be assigned to corresponding integral peaks integral values of *q*\*√2 and *√3. **H) Multiple Lorentz Function analysis of Braggs peaks from SAXS of standard LNP Particles**. Shown as black line are the SAXS data in a linear-linear plot. Multiple Lorentzian models were applied to deconvolute the data and are shown as dotted lines. The assignment of the H_||_ phase peaks with the spacing of d=54.6 Å and corresponding peaks at integral values of *q*\*√3 and *√4. The overlapping peak at *q*=0.127 Å^-1^ (d=49.5 Å) could be related to a corresponding integral peaks *q*\*√2, suggesting an additional cubosome phase. **I) SASVIEW Analysis**. Fitting of I vs q data shows core-shell sphere (light grey line) and core-shell ellipsoid model (dark grey lines) fits with polydispersity term. Residuals of the fits are shown in the lower panel. In this fitting, scattering length densities (SLDs) were fixed to calculated values and the shell diameter to that of a lipid bilayer (54 Å). The SLDs for the component parts of the DNA LNP formulations used in this fitting were calculated using MULCh. In the core-shell sphere model, a core radius of 101.8 Å ± 42.1 was determined χ^2^=2.7). In the core-shell ellispsoidal model fit, an equatorial core radius of 86.5 Å ± 86.9 and a long axis radius of 136.7 Å ± 332.5 was determined χ^2^=2.4) .

To further investigate the lipid organization and physical stability of CTS formulations, we performed a thermal analysis using differential scanning calorimetry (DSC) (**Fig. 2B**). This technique can identify phase transitions and structural ordering by measuring the energy required to heat samples, providing insights into molecular organization. Thermograms of standard LNPs displayed a single, broad, shallow exothermic peak centered around 80°C, suggesting a relatively amorphous structure with heterogeneous thermal transitions. In contrast, CTS LNPs yield two sharp, well-defined exothermic peaks: a major transition at approximately ∼75°C and a minor transition at ∼85°C. The presence of these distinct thermal transitions indicates that CTS LNPs possess a more ordered internal architecture with defined structural domains. The contrasting thermal profiles between the two formulations strongly suggest that the CTS protocol produces LNPs with a fundamentally different internal organization (supramolecular structure) compared to the standard formulation method.

We next applied different modalities of light scattering to quantitate the size and mass of these particles in solution. Using dynamic light scattering (DLS), we observe that CTS particles are almost two-fold larger in hydrodynamic radius (R_h_) in solution when compared to the standard LNP formulation, with a lower polydispersity index (PDI) (**Fig. 2C**). CTS particles had a determined R_h_ of 151 Å ± 2.3 (PDI = 0.14) compared to 80.2 Å ± 2.5 (PDI = 0.28) for the standard formulation. Consistent with these differences in size, the particles had very different retention times and weight-average mass profiles (M_w_), as determined by size-exclusion chromatography in-line with multiangle light scattering (SEC-MALS) (**Fig. 2D&E**). CTS particles had a M_w_ = 23.7 MDa ± 2.1% across the sizing peak, whereas standard particles were almost five-fold smaller in mass (M_w_ = 5.4 MDa ± 3.1%). Owing to the larger size of the CTS particles, we were also able to determine the radius of gyration (R_g_) for the CTS particles (38 nm ± 5.5%).

We next employed synchrotron small-angle X-ray scattering (SAXS) to further investigate the size and shape, and to better interrogate the internal structure of these particles. We employed in-line size-exclusion chromatography (SEC) to minimize and measure the possible effects of sample polydispersity in our analyses by performing singular value decomposition (SVD) analysis (see **Methods** and **Supplementary** Fig. 6). In this approach, very different scattering profiles were obtained for the CTS and standard particles (**Fig. 2F**), including well-defined primary Bragg peaks at *q*-values between ∼0.1-0.25 Å^-1^, indicative of a highly ordered internal structure (**Fig. 2G&H**). In this experimental configuration (where *q*_min_*D_max_ ≤ TT), it was possible to determine the size and shape of the standard LNP using the inverse Fourier transform (R_g_ of 156 Å, D_max_ of 400 Å), and to calculate *ab initio* electron density at low resolution (**Table 1** and **Supplementary** Fig. 6). It was not possible to apply the same conventional approaches to the CTS particles owing to their considerably larger sizes. However, empirical models for core-shell spheres and ellipsoids could be readily fit to the experimental data through the low and middle-q regime with a polydispersity term included, using calculated scattering length densities (SLDs) for the lipid and DNA components in X-rays and a fixed lipid bilayer depth of 54 Å. In the fitting, the resulting particle radii determined are largely consistent with the other measures reported herein (**Fig. 2I**). As seen in **Fig. 2G&H**, both samples show overlapping but distinguishable peaks. For the CTS particles, these are the peaks at *q*=0.10 Å⁻¹ and 0.12 Å⁻¹, while for the standard particles these are the peaks at q=0.115 Å⁻¹ and 0.125 Å⁻¹. Using the relationship *d*=2π/q, where *d* is the distance between the repeat lipid-DNA structures, the peaks indicated organized structure at *d* = 59.8 Å and 52.3 Å respectively for the CTS particle and 54.6 Å and 52.3 Å for the standard particles respectively (**Table 1**). The pattern of the Braggs peak in this data provides direct information on the structural arrangement through the reciprocal spacings between peaks. Inverse hexagonal phases (hexosomes) have peaks spaced in the pattern of √1, √3, √4.. and so on, while bicontinuous particles (cubosomes) have different integral spacings [22, 23] (√1, √2, √3..). In the CTS data, the *q*=0.10 Å^-1^ primary peak is readily assigned to a √1, √2, √3 spacing, indicative of cubosome phases (C), with corresponding peaks at *q*=0.15 Å^-1^ and 0.18 Å^-1^. The overlapping primary peak at *q*=0.12 Å^-1^ is readily assigned to corresponding peaks spaced in the pattern of √1, √2, √3, at *q*=0.18 Å^-1^ and 0.22 Å^-1^. Several additional weak peak features could not be assigned. In addition to primary peaks, a broader and less well-defined shoulder in the profile at *q*=0.06-0.09 Å^-1^ is observed which can be assigned to a higher order and more disordered phase. In contrast to the CTS structure, the standard LNP is better assigned with peak integrals consistent with an inverse hexagonal phase (H_||_), with the *q*=0.115 Å^-1^ peak readily assigned corresponding peaks at *q*=0.20 Å⁻¹ and 0.23 Å⁻¹. The overlapping peak at 0.125 Å^-1^ could be assigned as an additional cubosome phase with corresponding peaks at *q*=0.18 Å⁻¹ and 0.22 Å⁻¹, and like with the CTS particle, a region of disorder between *q*=0.05-0.1 Å^-1^ could also be assigned. These X-ray analyses indicate that the CTS particles feature a cubosome/cubic structure very distinct from the canonical inverse hexagonal phases that define the standard LNPs.

### CTS LNPs enhance DNA transfection by orders of magnitude

DNA-LNPs face five major “hurdles” to achieving high protein expression (**Fig. 3A**): 1) DNA structural perturbations during LNP synthesis, as conditions optimized for mRNA may destabilize DNA’s structure; 2) Upon cell uptake and endosomal escape, intracellular DNA sensors such as cGAS-STING repress protein translation; 3) Cytosolic DNases rapidly degrade DNA outside the nucleus; 4) DNA must navigate through the pores of the cytosolic gel; and 5) DNA must traverse the nuclear pore complex, which restricts passage of cargo >39 nm. We hypothesized that we could address the first hurdle by optimizing synthesis conditions to preserve DNA structure. We further hypothesized that we could address the next four hurdles by *compactifying* DNA into small spheres (“cores”), which would protect the DNA from DNA sensors and DNases (hurdles 2 & 3) and enable easier transit of DNA through the narrow passages of the cytosolic gel and nuclear pore complex (hurdles 4 & 5). More specifically, we hypothesized that CTS could produce DNA cores that possessed such properties.

**Figure 3:**
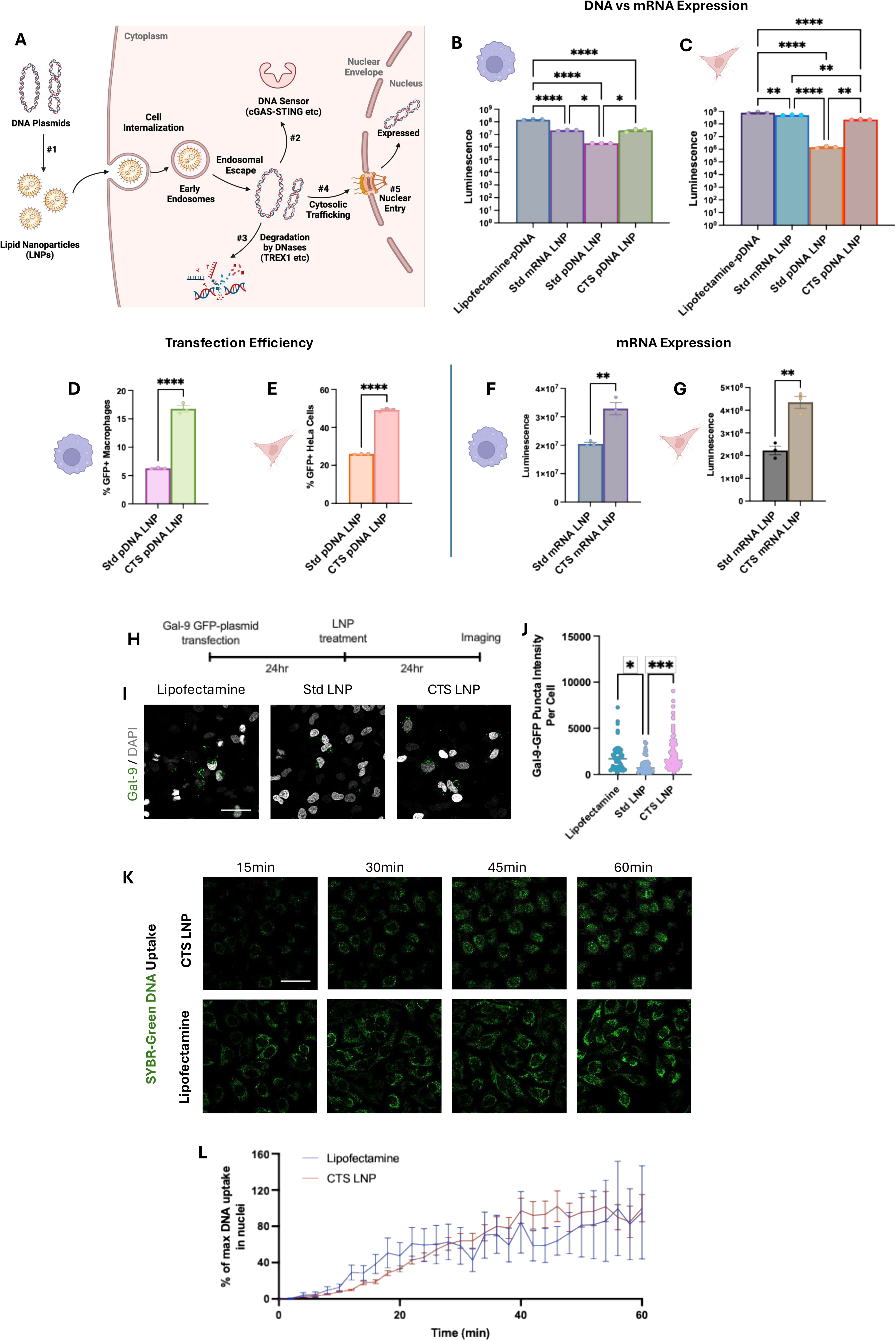
**Core-then-shell LNPs enhance transfection by orders of magnitude via improving endosomal escape and rapidly localizing DNA to the nucleus. A) DNA LNPs have 5 major hurdles to high protein expression**: DNA structural perturbations during the LNP synthesis (#1); Upon cell uptake and endosomal escape, the DNA can be recognized by intracellular DNA sensors such as cGAS-STING (#2) and cleaved by DNases (#3); Finally, the DNA must travel through the cytosol (#4) and cross into the nucleus to engage in transcription (#5). The current study aims to employ structural modifications of DNA LNPs to overcome these hurdles. B-G) *In vitro* transfection with the listed LNPs in RAW macrophages (B, D, F) and HeLa epithelial cells (C, E, G) shows that comparing with lipofectamine, a gold standard benchmark for *in vitro* transfection. CTS DNA LNPs express 10-fold higher than standard DNA LNPs in RAW macrophages and 100-fold higher in HeLa cells, and equivalent to mRNA-LNPs. In addition, CTS LNP has significant transfection efficiency than Std LNP. Not only for DNA, CTS mRNA LNPs also enhance mRNA expression, compared to standard mRNA-LNPs. **H) Endosome escape events.** To measure endosomal escape events, HeLa cells were transfected with plasmids containing GFP-labeled galectin-9, which forms puncta around endosomes that have opened. **I) Representative image of Gal-9 reporter cells after LNP or lipofectamine treatment. J) Quantification of Gal-9 puncta.** The Gal-9-GFP puncta per cell were higher in CTS LNP compared to standard LNPs, and reached a similar level to the gold- standard Lipofectamine. **K) Representative image of time lapse of DNA uptake in HeLa cells. L) Quantification of DNA uptake.** To measure DNA uptake and import into the nucleus, DNA was fluorescently labeled with SYBR-Green, loaded into CTS-LNPs, and given to HeLa cells. The uptake into the nucleus via CTS LNPs is on a very similar time course to that of Lipofectamine. Scale bar denotes 50 µm.

We began by focusing on hurdle #1 above. The standard synthesis conditions of LNPs may alter DNA’s structure in deleterious ways. In mRNA-LNP synthesis, the nucleic acid is dissolved in pH 4 citrate buffer to maximize protonation of ionizable lipids [2]. However, this low pH may destabilize the DNA helix and cause conformational changes from the biologically active B-form, the most common biologically active structure present in cells under normal physiological conditions. Other conformations can include A-form, Z-form, G-quadruplex, and others, as well as less defined conformations that might occur after partial acid hydrolysis of the DNA [24].

To assess the effect of buffer and pH on DNA conformations, we analyzed DNA in different buffers by circular dichroism (CD) spectroscopy, which measures B-form DNA via relative magnitude of the ellipticity signal in the range of 200 - 320 nm. A typical B-form DNA has a positive maximum around 290 nm, a negative maximum around 245 nm and a broad positive peak between 260 - 280 nm. We compared pH 4 citrate (classic mRNA-LNP synthesis buffer) vs pH 6 TBS with 4 mM Mg2^+^. Tris is the most common buffer used in DNA reactions, and Mg^2+^ stabilizes the B-form DNA helix via a variety of mechanisms, mostly involving electrostatic interaction between Mg^2+^ and DNA’s phosphates [25]. As shown in **Supplementary** Fig. 4A, the CD spectroscopic signal for B-form DNA is significantly disrupted by citrate buffer vs TBS buffer. Titrating Mg2+ in the TBS buffer, we found that 4 mM Mg2+ slightly improves B-form DNA signal in CD spectroscopy, compared to 0 and 12.5 mM Mg2+ (**Supplementary** Fig. 4B).

Having established the impact of buffer conditions on DNA conformation, we next evaluated how these conditions affect DNA-LNPs’ transfection efficiency. We synthesized luciferase-encoding DNA-LNPs in different buffers, dialyze against buffers containing corresponding Mg^2+^ as in their DNA dissolving buffer, and tested their performance in RAW 264.7 macrophages and HeLa cells (**Supplementary** Fig. 5). The Tris-based buffers (TAE) at pH 5 outperformed citrate buffer (pH 4). Critically, we discovered that Mg^2+^ in the dialysis buffer was essential for expression, likely due to its role in stabilizing DNA structure - though notably, this benefit was only observed at pH > 4, as Mg^2+^ showed no positive effects in acidic conditions (**Supplementary** Fig. 6), possibly due to weakened Mg^2+^-DNA interactions in proton-rich environments [26]. A comprehensive screen of buffer conditions across both cell types revealed that Tris-based buffers at pH 6 with Mg^2+^ consistently produced the highest expression levels, with TBS showing particularly strong performance in macrophages and TAE in HeLa cells. Cryo-EM analysis confirmed that these buffer-optimized LNPs (TBS, pH 6, Mg^2+^) maintained similar morphology to standard LNPs while achieving significantly higher expression levels (**Supplementary** Fig. 2C).

With optimized buffer conditions maintaining DNA structure during synthesis, we next investigated how CTS architecture could overcome the remaining intracellular hurdles, all of which were hypothesized to benefit from the DNA compactification of CTS. We compared luciferase expression from plasmid DNA delivered via CTS LNPs vs standard DNA-LNPs. CTS LNPs yield >10x higher expression than standard LNPs in RAW macrophages, and ∼150-fold higher in HeLa cells, with the levels comparable to mRNA-LNPs (**Fig 3B-C**). Comparing transfection efficiency, quantified as % of cells that are transfected by plasmid encoding GFP, CTS DNA-LNPs outperform standard LNPs by three-fold in RAW macrophages and two-fold in HeLa cells (**Fig 3D-E**). Thus, compared to standard LNP formulations and single-step mixing, using MSM to create CTS LNPs improves protein expression from DNA cargo by 1-3 orders of magnitude.

We had originally hypothesized that the CTS process would primarily work via benefits of compact DNA packaging, enhancing cytoplasmic trafficking and nuclear entry. However, another possibility is that CTS LNPs might affect LNPs’ endosomal escape, which is typically the rate limiter for LNPs’ transfection of mRNA [27]. To test this, we applied the same MSM CTS design for mRNA delivery and tested the protein expression level of CTS-mRNA-LNPs in both RAW and HeLa cells, figuring out that the augmentation of expression of CTS-mRNA-LNPs is 2-4 fold, compared to standard mRNA-LNP (**Fig. 3F-G**). The enhanced mRNA expression indicates that CTS design alleviates the hurdles imposed by endo/lysosomes including inefficient endosomal escape. However, the expression improvement in the mRNA system is much less than the multiple orders of magnitude improvement for DNA. This could give real mechanistic insight: the bulk of CTS’s benefits for DNA delivery is DNA-specific. This lowers the probability that CTS predominantly improves DNA transfection via effects on endosomal escape.

To quantify endosomal escape events, we transfected HeLa cells with a galectin (Gal)-9- GFP plasmid and monitored LNP-mediated endosomal disruption (**Fig. 3H**). Galectins are primarily expressed in the cytosol and can be recruited to endosomes when membrane are damaged. Under baseline conditions, cells showed diffuse cytoplasmic Gal-9-GFP distribution [28]. Upon treatment with different LNP formulations, we observed distinct Gal-9-GFP puncta formation, indicating sites of endosomal membrane disruption (**Fig. 3I**). Quantification revealed that CTS LNPs induced higher Gal-9-GFP puncta formation than standard LNPs (**Fig. 3J**). However, this modest (though statistically significant) enhancement in endosomal escape cannot fully account for the 1-3 orders of magnitude improvement in transfection observed with CTS LNPs.

We next examined nuclear delivery kinetics using fluorescently labeled DNA. Live-cell imaging in HeLa cells revealed that CTS LNPs achieved nuclear accumulation rates comparable to Lipofectamine over a 60-minute period (**Fig. 3K**). Quantitative analysis showed nearly identical uptake profiles for both delivery systems (**Fig. 3L**), suggesting that while CTS LNPs match the nuclear delivery efficiency of this gold-standard transfection reagent, additional mechanisms beyond endosomal escape and nuclear entry must contribute to its superior transfection capabilities.

### Co-loading CTS DNA-LNPs with STING inhibitors further increases expression and enables in vivo expression peaks comparable to mRNA-LNPs

Before moving CTS DNA-LNPs *in vivo*, it was essential to address whether their DNA cargo activates intracellular DNA-sensors, especially cGAS-STING. Our previous work has shown that standard DNA-LNPs strongly activate STING *in vivo*, leading to significant toxicity upon intravenous administration [13], and represses translation [29].

We had initially hypothesized that the CTS structure would provide enhanced protection of DNA through compaction in the core structure, shielding it from both cGAS-STING sensing and DNase degradation after endosomal escape (**Fig. 3A**). However, our findings indicate CTS LNPs’ improved endosomal escape (**Fig 3H-J**) may actually increase cytosolic DNA accumulation over time, enhancing activation of DNA sensors like cGAS-STING. To test this, we incubated RAW macrophages with CTS DNA-LNPs, standard DNA-LNPs, or Lipofectamine with DNA and measured STING activation by immunostaining of phosphorylated STING [30]. We found that CTS LNPs indeed activated STING more strongly than standard LNPs, reaching levels comparable to Lipofectamine-based DNA delivery (**Fig. 4A-B**).

**Figure 4:**
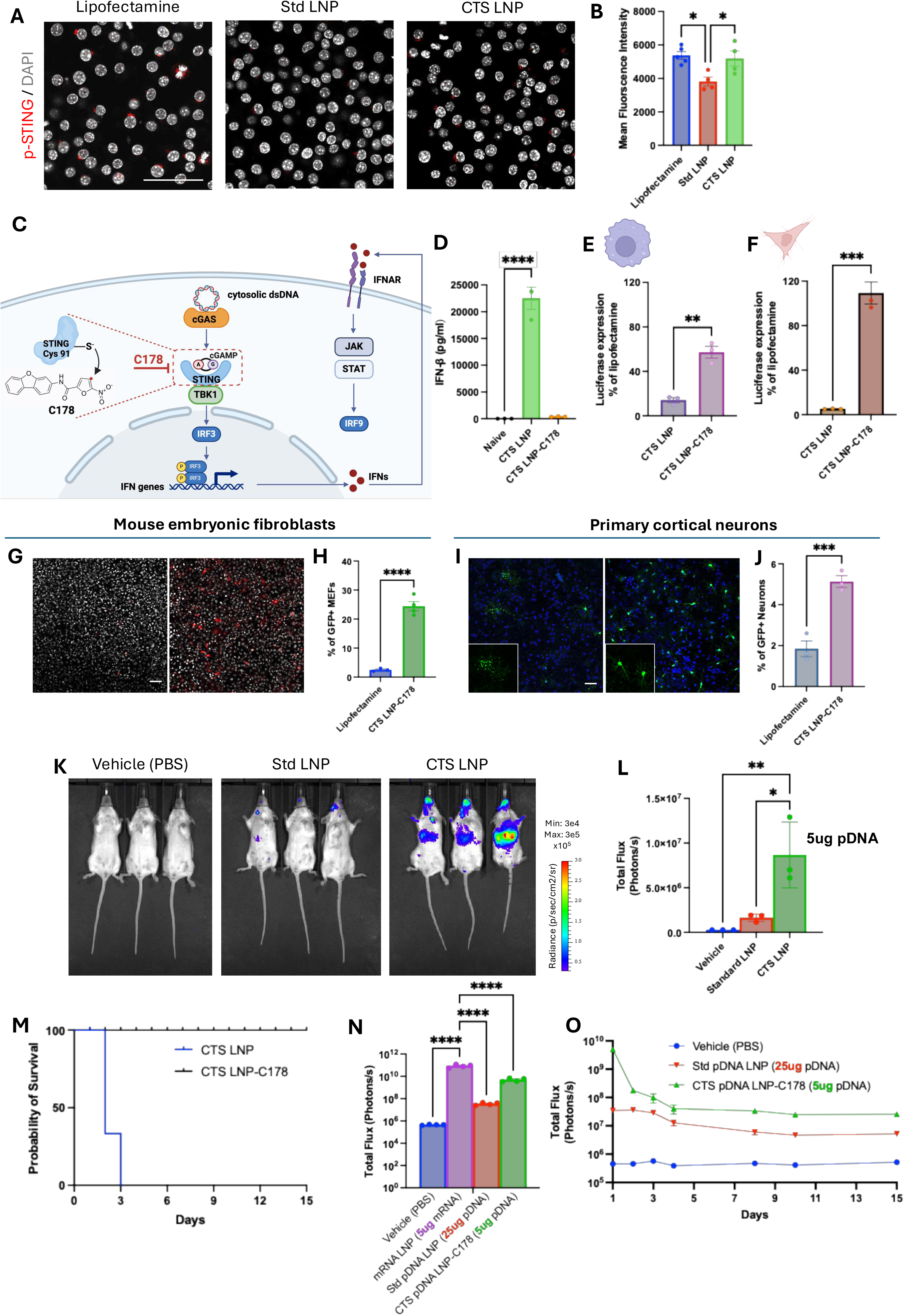
Co-loading a STING-inhibitor into core-then-shell (CTS) DNA LNPs enables high-level, long-term expression of DNA LNPs in primary cells in culture and *in vivo*. A) Representative image of phosphate-STING (p-STING) signal in RAW macrophages after LNP or lipofectamine treatment. The cGAS-STING pathway detects cytosolic DNA and thereafter activates pathways involved in inflammation and repression of protein translation. To measure STING activity, we stained RAW macrophages with phospho-STING antibody. **B) Quantification of p-STING.** We found that CTS LNPs activate STING much more than standard LNPs, and to the same extent as Lipofectamine. **C) Schematic illustration of cGAS-STING activation pathway and C-178 function in inhibition of STING.** STING induces expression of IFN- β, which activates inflammation and represses translation. C-178 is a small molecule analog of the endogenous negative feedback on STING. **D) IFN- β level.** CTS DNA LNPs massively increase expression of IFN-β by RAW macrophages, but this is completely inhibited by co-loading C-178 into the CTS LNPs. **E, F) *In vitro* transfection of CTS LNP co-loading with C-178.** In both RAW macrophages and HeLa epithelial cells, C-178 co-loading improves protein expression. **G, H, I, J) Primary cell transfection by CTS LNPs.** Lipofectamine is the gold standard for *in vitro* transfection, but is inefficient with most primary cells. Here we measured expression, via GFP plasmids, in mouse embryonic fibroblasts (G, H), and mouse primary cortical neurons (I, J). In both cases, CTS LNPs co-loaded with C-178 had markedly higher transfection efficiency than Lipofectamine, and preserved neuron morphology. **K) Representative images from In Vivo Imaging System (IVIS).** Standard and CTS LNPs, without any STING inhibitor (e.g., C-178), were IV-injected into mice and luciferase expression was measured via IVIS at 24 hours. **L) IVIS image quantification.** CTS showed expression markedly above standard LNPs, but the CTS s’ expression was likely inhibited by STING expression. **M) Animal survival.** CTS LNPs without STING inhibition had 100% mortality within 3 days, but CTS LNPs co- loaded with C-178 had 0% mortality. **N) Day 1 IVIS image quantification.** After IV injection, standard DNA LNPs drive expression that is several orders of magnitude lower than mRNA LNPs, though we recently showed that these DNA LNPs drive expression for more than 3 months, while LNP-delivered mRNA decays in days. However, CTS DNA LNPs with co-loaded C-178 show peak expression levels orders of magnitude higher than standard-DNA-LNPs, with the dose 5 times lower than standard-DNA-LNPs, and comparable to mRNA s’ peak. **O) Longitudinal quantification of DNA expression *in vivo.*** Scale bar denotes 50 µm.

To mitigate STING activation, we loaded our CTS DNA-LNPs with C-178, a small molecule STING inhibitor that covalently binds to Cys91, which is the same mechanism employed by mammal’s endogenous STING inhibitors, nitrated lipids (**Fig. 4C**) [31]. C-178 was efficiently incorporated during CTS LNP synthesis, with a drug entrapment efficiency of 97.5% and drug-to-lipid molar ratio of 0.025. The loading of C-178 did not affect DNA entrapment efficiency. CTS LNPs loaded with C-178 effectively suppressed STING-mediated immune responses, as evidenced by complete elimination of IFN-β production in RAW macrophages (**Fig. 4D**). Importantly, C-178 co-loading enhanced transgene expression by approximately 4-fold in RAW macrophages and 20-fold in HeLa cells, compared to CTS LNP only (**Fig. 4E-F**).

Having solved the STING activation issue, we next evaluated STING-inhibiting C-178- loaded CTS DNA-LNPs in challenging primary cell types. In mouse embryonic fibroblasts, C-178-loaded CTS DNA-LNPs significantly outperformed Lipofectamine in transfection efficiency (**Fig. 4G-H**). Similarly, in primary cortical neurons, traditionally considered highly resistant to transfection, our formulation achieved superior gene delivery compared to Lipofectamine and preserved neuron morphology (**Fig. 4I-J**).

*In vivo* studies demonstrate that CTS DNA-LNPs without C-178 enhance liver luciferase expression 5-fold over standard DNA-LNPs (**Fig. 4K-L**), but also induce significant toxicity leading to mortality (**Fig. 4M**). Loading CTS LNPs with C-178 eliminates this toxicity while maintaining high transgene expression. A 5 ug dose of plasmid DNA in CTS DNA-LNPs loaded with C-178 achieved orders of magnitude superior expression to a 25 ug dose in standard LNPs – despite the 5-fold reduction in dose. C-178-loaded CTS DNA-LNPs produced peak expression levels approaching those of mRNA-LNPs (**Fig. 4N**), but with the crucial advantage of sustained expression (**Fig. 4O**). The ability to achieve mRNA-like expression peaks with a reduced DNA dose, while maintaining long-term expression, makes CTS DNA-LNPs very promising for treating chronic diseases.

## DISCUSSION

These studies show that LNPs’ supramolecular structure (spatial arrangement of molecules) has a major effect on their performance. Further, we show that supramolecular structure can be controlled not just by the constituent molecules’ identity, but also by the methodology of mixing. In particular, multi-stage mixing (MSM) allows for control of supramolecular structure, even enabling capture of favƒorable metastable states, while maintaining continuous flow production. MSM’s simplest protocol, a core-then-shell (CTS) two-layer synthesis, improved numerous LNP performance metrics, including: lower polydispersity index (PDI) (**Fig. 1E**); a much lower fraction of “empty” LNPs (undetectably low concentration of nucleic acid cargo) (**Fig. 1N**), which we previously showed are inflammatory; prevention of side effects (e.g., clotting) by burying particular lipids in the core (**Supplementary** Fig. 1); improved protein expression by mRNA-LNPs (**Fig. 3F-G**); and most impressively, improvement in DNA-LNPs’ protein expression by 2-3 orders, bringing it to levels similar to the peak of mRNA-LNPs (**Figs. 3 & 4**).

This enormous improvement in DNA-LNPs’ expression may now allow DNA-LNPs to serve a long-standing goal in genetic medicine: common chronic diseases. These diseases include the top causes of death like atherosclerosis, heart failure, and chronic obstructive pulmonary disease (COPD), as well as the top causes of illness and disability, such as Alzheimer’s disease, chronic pain, and chronic kidney disease [32]. Unfortunately, genetic medicine tools have thus far been best suited to either rare monogenic diseases (AAV and CRISPR) or vaccines (mRNA-LNPs). By contrast, DNA-LNPs offer unique advantages that make them particularly well-suited for treating common chronic diseases: they can express or knockdown (by expressing short hairpin RNA [shRNA]) even large cargo genes because they impose no major cargo capacity constraint (unlike AAV’s rigid capsid) [33]; maintain expression for ∼6 months per dose (unlike the hours of expression with mRNA-LNPs) [32]; achieve genetically-encoded cell-type specificity with inclusion of large promoter regions (not possible with mRNA-LNPs); avoid permanent integration or disruption of the genome (unlike AAV and CRISPR) [34]; minimize immunogenicity (unlike AAV) [34]; and leverage existing mRNA-LNP targeting technologies. Thus, DNA-LNPs can, with continued development, fill the greatest gap in genetic medicine, opening up therapy for dozens of common chronic diseases.

Importantly, the CTS LNPs developed here represent a fundamentally different approach from traditional core-shell nanoparticles created through layer-by-layer (LbL) techniques. While LbL methods typically involve sequential deposition of oppositely charged materials onto a preformed core [9], our multi-stage mixing (MSM) approach employs a sophisticated microfluidic design that achieves two critical innovations simultaneously: (1) controlled compact packaging of DNA using precisely titrated DNA-condensing agents that facilitate nuclear entry; (2) rapid capture of thermodynamically metastable core matrices by lipid shells before aggregation can occur. The microfluidic architecture is crucial to this process. Traditional LbL requires multiple separation and purification steps between layer additions, which can lead to material loss and batch-to-batch variability [9]. In contrast, our MSM system achieves precise spatiotemporal control over the mixing process in a continuous flow format. The first mixing stage creates DNA-lipid complexes with moderate compacting - critically maintaining an N:P ratio below 2 to avoid over-condensation that would inhibit nuclear entry [17]. The second stage rapidly envelopes these nascent cores with a lipid shell before they can grow beyond the optimal size range. This process occurs in milliseconds, preventing the formation of large aggregates (>2000 nm) that typically plague attempts to create DNA-lipid cores through bulk mixing methods.

Our MSM approach offers distinct advantages over conventional core-shell particle synthesis through its precise control over the stoichiometry of the compact lipoplex DNA core, allowing optimization for nuclear transport. The system enables capture of metastable intermediates that would be impossible to achieve through equilibrium processes, while maintaining continuous flow production that enhances reproducibility and scalability. Compared to multi-step LbL processes, our method minimizes material loss and prevents aggregation through rapid shell formation.

While we have shown the CTS structure massively improves transfection, there is great need for future work deeply exploring the mechanisms of CTS’s improved efficacy. We were guided in our design of CTS by the hypothesis that compactifying DNA into a core would protect it from intracellular DNases, and enable it to pass more easily through the viscous cytosol and the narrow nuclear pore complex (**Fig. 3A**). However, proving whether or not these are indeed the mechanisms of CTS’s improved efficacy will require additional studies. Thus far, we have a few glimpses at the mechanisms. First, we found that it is essential to maintain DNA in the B-form, by optimizing the buffers in which DNA is formulated into LNPs and into which LNPs are dialyzed following synthesis. Second, we found that CTS improves mRNA-LNPs’ transfection, but only by ∼2-4-fold, which is much less than the ∼1,000-fold improvement by CTS for DNA-LNP’s transfection. Third, we found that CTS improves endosomal escape (measured by Gal-9-GFP puncta), but only by <2-fold. Fourth, we found that the time course of nuclear translocation of DNA is similar for CTS DNA-LNPs and Lipofectamine. These four observations allow us to roughly estimate the relative contributions of different mechanisms to CTS’s improved transfection: maintenance of the B-form plays a moderate role; improved endosomal escape plays a modest role (based on Gal-9 and mRNA expression data); and CTS allows rapid translocation to the nucleus.

While these tentative conclusions on mechanism are consistent with our initial hypothesis, we need more definitive mechanistic studies in the future. Such studies could include comparing CTS vs standard DNA-LNPs in the following ways: measuring degradation of cargo-DNA after it enters the cell, including whether the DNA is partially damaged (nicked, linearized, or conformational changes); tracking the movement of cargo-DNA through the cytosol and nuclear pore complex, especially after plasma membrane permeabilization (to eliminate the effect on endosomal escape); and modifying CTS to produce larger or smaller cores, and compactify the DNA with different cations or other species, to probe structure-function relationships. Such mechanistic studies will not only elucidate key science in genetic medicine and material science, but also guide further practical engineering of MSM and CTS LNPs.

Indeed, MSM and CTS LNPs may open up a vast design space of materials for genetic medicine. Customized MSM designs may allow for multiple shell layers. Within just the CTS architecture, there is tremendous room for optimization, possibly allowing loading of chemical species that would normally partition out of LNPs. For example, we can change out the molecules used to condense the DNA and vary the microfluidic design to change the size and material properties of the LNPs. Finally, there is exciting work to be done to scale-up the manufacturing of MSM and CTS, which may be done with highly parallelized microfluidics [35, 36], or perhaps with macro-scale mixers like the confined impinging jet mixers used for the COVID vaccines [7].

With continued innovation of MSM and CTS LNPs, it is possible to foresee a time when DNA-LNPs achieve their promise of safe, long-term control of expression of any protein. Such DNA-LNPs would be able to treat a vast number of common chronic diseases. All of this can be made possible by applying novel materials engineering techniques like MSM to control the supramolecular structure of LNPs.

## MATERIALS AND METHODS

### Material

SM-102 was purchased from Echelon Biosciences (Ontario, Canada). 18:0 PC (DSPC, 1,2-distearoyl-sn-glycero-3-phosphocholine), DMG-PEG 2000 (1,2-dimyristoyl-rac-glycero-3- methoxypolyethylene glycol-2000), 18:1 (Δ9-Cis) PE (DOPE), 18:1 TAP (DOTAP), 18:0 PE-DTPA and cholesterol were purchased from Avanti Polar Lipids (Alabaster, AL). RAW 264.7, HeLa cells, and mouse embryonic fibroblasts (MEFs) were purchased from American Type Culture Collection (Manassas, VA). Reporter lysis buffer and luciferin assay solution were purchased from Promega (Madison, WI). Plasmid DNA (pDNA) was purchased from Aldevron (<0.1 EU/µl endotoxin level). 5moU nucleoside-modified firefly luciferase mRNA was purchased from TriLink BioTechnologies. Indium-111 chloride (In-111) was purchased from BWXT Medical (Ottawa, Canada). Sodium iodine (I-125) was purchased from PerkinElmer (Waltham, MA). All other chemicals and reagents were purchased from Fisher Scientific (Hampton, NH), unless otherwise noted.

### Animals

All animal studies were carried out in accordance with the Guide for Care and Use of Laboratory Animals under the protocol approved by the University of Pennsylvania Institutional Animal Care and Use Committee, and conformed to all relevant regulatory standards. Naive C57BL/6 mice (male, aged 6-8 weeks, 23-25g), naive BALB/c mice (female, aged 6-8 weeks, 23- 25g) were procured from the Jackson Laboratory (Bar Harbor, ME). The animals were housed in a controlled environment maintained at temperatures between 22–26 °C, with a 12/12-hour light/dark cycle, and provided with access to food and water.

### Cell culture

RAW264.7 mouse macrophages were cultured in Dulbecco’s modified Eagle’s medium (DMEM) with 10% heat-inactivated fetal bovine serum (FBS) and 1% penicillin/streptomycin (PS). HeLa cells were cultured in Eagle’s Minimum Essential Medium (EMEM) supplemented with 10% FBS and 1% PS. Mouse embryonic fibroblasts (MEFs) were cultured in DMEM supplemented with 10% FBS, 2 mM L-GlutaMAX, 1X non-essential amino acids, 1 mM sodium pyruvate, 10 mM HEPES, 100 U/mL penicillin, 100 mg/mL streptomycin. All cells were incubated with 5% CO_2_ at 37°C.

### LNP Synthesis

LNPs were formulated using NanoAssemblr Ignite nanoparticle formulation systems (Precision Nanosystems, Vancouver, Canada), or using the microfluidics device designed in-house (**Fig. 1C**). The aqueous phase was made in different buffers including 50 mM citrate buffer (pH 4.0), Tris-acetate-EDTA (TAE), Tris-buffered saline (TBS), and HEPES with pDNA. For standard LNPs, an organic phase containing a mixture of lipids dissolved in ethanol at a designate molar ratio (**Supplementary** Fig. 2A) was mixed with the aqueous phase containing DNA, at a flow rate ratio of 1:3 and at a total lipid to nucleic acid weight:weight ratio of 40:1. For CTS LNPs, channel 1 (core lipid phase) contains lipid mixture of DOPE and DOTAP; channel 2 (shell lipid phase) contains lipid mixture of ionizable lipid, DSPC, PEGylated lipid and cholesterol at a designated molar ratio (**Fig. 1A**); channel 3 (core aqueous phase) is pDNA in aqueous buffer. 3 channels were mixed at a flow rate ratio of 1:1:3. For drug-loaded CTS LNPs, STING inhibitor C- 178 was added to channel 2.

LNPs were dialyzed against 1X PBS or 1X PBS containing MgCl_2_ for 2 hours. Mg^2+^ concentration in the dialysis buffer corresponds to its concentration in the DNA-containing aqueous buffer, unless otherwise stated. After dialysis, LNPs were sterilized using a 0.22 µm filter, and stored at 4 °C for later use.

### In-House Microfluidics Device Fabrication

A custom, dual-stage microfluidic mixer was fabricated in the Quattrone Nanofabrication Facility at the University of Pennsylvania. Design of the device was completed in AutoCAD (Autodesk, San Rafael, CA). Briefly, using a mask aligner, each layer of the multilayer architecture was photolithographically patterned onto a 100mm silicon wafer (ID 775; University Wafer, South Boston, MA) spray coated with S1805 photoresist (Dow, Midland, MI). After patterning and development, deep reactive ion etching (SPTS Rapier Si DRIE, Newport, UK) was performed. After the silicon was patterned and etched, the device was anodically bonded to a 100-mm Borofloat 33 glass wafer (ID 517; University Wafer) using an EVG 510 Wafer Bonding System (EVG Group, Oberosterreich, Austria).

Dual-valve pressure regulators (Alicat Scientific, Tucson, AZ) were used to pressurize 15mL vessels (Elveflow, Paris, France), with outputs mechanically coupled to the input ports of the microfluidic mixer. Separate inputs were provided for the ethanol and aqueous phase components, to allow for a two-step nanoprecipitation. The micromixer accommodated individual mixing of constituent components of the ethanol and aqueous phases to separately form the core and shell, which then combined and underwent further mixing. Pressures were chosen to allow the total flow rate through each input to combine to a total flow rate of 1.2mL/min. CTS LNP samples were collected directly into a dialysis cassette (Thermo Fisher, Waltham, MA), and dialyzed against 1X PBS containing 4 mM MgCl_2_ for 2 hours.

### LNP Characterization

Hydrodynamic nanoparticle size, polydispersity index (PDI), and bulk zeta potential of LNPs were measured by dynamic light scattering using Zetasizer Pro ZS (Malvern Panalytical, Westborough, MA). Distribution of zeta potential of individual LNP was measured by ZetaView instrument by Particle Metrix (Inning am Ammersee, Germany).

Encapsulation efficiency and concentration of pDNA or mRNA were measured using a Quant-iT-PicoGreen, or Ribogreen, respectively. 50 μL of TE buffer and 50 μL of 2% Triton X-100 (in TE buffer) were added in duplicates to a black bottom 96-well plate. LNP formulations were diluted in the TE buffer, and 50 μL of each formulation was added to the TE buffer and 2% Triton X-100. After 10 minutes of incubation time, RiboGreen or PicoGreen reagent was diluted 1:100 or 1:200, respectively, in the TE buffer and added to each well. Picogreen fluorescence was measured at excitation of 480 nm and emission at 520 nm, and RiboGreen signal was measure using an excitation 485 nm of and emission of 528 nm by a plate reader (Spectramax M2; Molecular Devices, San Jose, CA).

Drug encapsulation was measured by high-performance liquid chromatography, and drug entrapment efficiency was calculated as weight of loaded drug in LNPs over weight of total drug added into LNP formulation.

### Cryo Electron Microscopy

To prepare samples for cryo-EM imaging, lacey carbon–coated 300 mesh copper grids (Electron Microscopy Sciences) were glow discharged at 15 mA for 30 s with the PELCO easiGLOW glow discharge system (Ted Pella). Then, 8 μL of sample was applied to the grids. The grids were blotted with filter paper and plunge frozen in liquid ethane using Vitrobot Mark IV, under 4 °C and 100% humidity. The grids were kept in liquid nitrogen until imaging. Cryo-EM images were collected with Titan Krios Cryo-TEM (ThermoFisher).

### Circular Dichroism Spectroscopy

All CD experiments were carried out using Jasco 1500 CD spectrometer. The following parameters were used: range 200-320 nm, data pitch = 1 nm, CD scale = 200 mdeg/0.1 dOD, FL scale = 200 mdeg/0.1 dOD, Digital Integration Time = 1 sec, bandwidth = 1 nm. pDNA were measured at the concentration of 30 µg / mL. Buffer solutions alone were used as blanks to establish baselines for correction. Normalized raw CD data were used for plotting and analysis.

### Differential Scanning Calorimetry

Samples of LNPs were prepared for calorimetry measurements at particle concentrations of 3-5e11 nanoparticles per mL in PBS. PBS was used as a reference solution for all differential scanning calorimetry measurements. Reference PBS was drawn from the same aliquot used to dilute nanoparticle samples. Lipid nanoparticle suspensions and reference solutions were heated from 4°C to 130°C at a target rate of 1°C per minute in a TA Instruments Nano DSC operating in scanning mode. Background measurements of reference PBS vs. reference PBS were obtained to confirm stability of the measurements and reference vs. reference thermograms were subtracted from all LNPs vs. PBS reference thermograms. Sigmoidal fits were applied to baselines of background-corrected thermograms in TA Instruments NanoAnalyze software. After baseline subtraction, thermogram data were normalized to instantaneous scan rate and molar concentration of nanoparticles as measured by nanoparticle tracking analysis. The resultant molar heat capacity thermograms were integrated to yield enthalpy data or normalized to temperature then integrated to yield entropy data. Enthalpy, entropy, and Gibbs energy of any phase transitions were extracted from this data.

### Small Angle X-ray Scattering

#### Dynamic Light Scattering

A Nanobrook Omni particle sizer (Brookhaven Instruments Corporation, Holtsville, NY, USA) was used to record data at 25°C in polystyrene 1-cm cells using a standard diode laser at 640 nm, with scattering recorded at an angle of 90°. Three scans were recorded for each sample and hydrodynamic radii (R_h_) were calculated using the BIC Particle Solutions software v3.6.0.7122.

#### SEC-MALS

100 μL of DNA-LNP at a particle concentration of ∼10^11^ particles were injected and eluted isocratically at 0.2 ml/min from a 7.8 mm TSKgel G6000PWxl-CP sizing column equilibrated in 1X PBS at room temperature. Absolute molar mass of the DNA-LNPS was determined in-line using multi-angle light scattering. Light scattering from the column eluent was recorded at 18 different angles using a DAWN-HELEOSII MALS detector (Wyatt Technology Corp.) operating at 658 nm. Eluent concentration using a mass averaged dn/dc value of 0.16 cm3/g was determined using an in-line Optilab T-rEX Interferometric Refractometer (Wyatt Technology Corp.). The weight-averaged molar masses of species within defined chromatographic peaks were calculated using the ASTRA software version 8.2.2 (Wyatt Technology Corp.), by construction of Debye plots (KC/Rθ versus. sin^2^[θ/2]) at one second data intervals. The weight-averaged molar mass was then calculated at each point of the chromatographic trace from the Debye plot intercept and an overall average molar mass was calculated by weighted averaging across the peak. Normalization coefficients for diode counting efficiency were obtained using a BSA standard.

#### SEC-SAXS

*Synchrotron Size-Exclusion Chromatography in line with Small Angle X-ray Scattering*. Size-exclusion chromatography (SEC)-SAXS data were collected at beamline 16-ID (LiX) of the National Synchrotron Light Source II (Upton, NY) [37]. Data were collected at a wavelength of 1.0 Å in a three-camera configuration, yielded accessible scattering angle where 0.006 < q < 3.0 Å^-1^, where q is the momentum transfer, defined as q = 4π sin(θ)/λ, where λ is the X-ray wavelength and 2θ is the scattering angle; data to q<0.5 Å^-1^ were used in subsequent analyses. 100 μL of DNA-LNP at a particle concentration of ∼10^11^ particles were injected and eluted isocratically at 0.2 ml/min from a 7.8 mm TSKgel G6000PWxl-CP sizing column equilibrated in 1X PBS (Invitrogen), at room temperature. Eluent from the column flowed into a 1 mm capillary for subsequent X-ray exposures at 1-s intervals. Plots of intensity from the forward scatter closely correlated to in-line UV and refractive index (RI) measurements.

#### SAXS Analysis

SEC-SAXS data sets were analyzed in the program RAW [38]. Buffer subtracted profiles were analyzed by singular value decomposition (SVD) and the ranges of overlapping peak data determined using evolving factor analysis (EFA) as implemented in REGALS [39]. The determined peak windows were used to identify the basis vectors for each component and the corresponding SAXS profiles were calculated. When manually fitting the pair distribution function P(r) using the program DIFT[40], the maximum diameter of the particle (D_max_) was incrementally adjusted to optimize χ^2^ figures, to minimize the discrepancy between the fit and the experimental data to a *q_max_* ≤ 8/R_g_, and to optimize the visual qualities of the distribution profile. Deconvolution of the primary Braggs peaks in SAXS were performed using multiple Lorentz with the built-in functions implemented in OriginPro 2024b (Northampton, MA, USA), as previously described [23]:

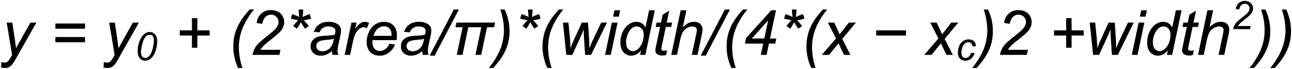

where y_0_ is an offset and was set to the X-ray baseline. x_c_ is a center of function and corresponds to the center of the SAXS peak. In the fitting, the peak position was fixed and the area and width of the peaks were not restrained. Peaks were assigned to inverse hexagonal (H_||_) or lamellar (L_α_) phases based on peaks corresponding to the first order Bragg peak and corresponding peaks at integral increments [22].

SASView (www.sasview.org) was used to fit experimental data to empirical models for core-shell sphere and ellipsoid models. In this fitting, scattering length densities (SLDs) were fixed to calculated values and the shell diameter to that of a lipid bilayer (54 Å). The SLDs for the component parts of the DNA-LNP formulations used in this fitting were calculated using MULCh [41].

DENSS [42] was used to calculate the *ab initio* electron density map directly from the DIFT output. Twenty reconstructions of electron density were performed in the slow mode with default parameters and subsequently averaged and refined. Reconstructions were visualized using PyMOL 2.5.2 Molecular Graphics System (Schrodinger, LLC, New Your, NY, USA) with five contour levels of density rendered with these respective colors: 15σ (red), 10σ (green), 5σ (cyan), 2.5σ (blue), and -0.7σ (blue). The sigma (σ) level denotes the standard deviation above the average electron density value of the generated model.

### *In vitro* LNP transfection

HeLa cells and RAW 264.7 macrophages were seeded in 96-well plates and incubated overnight (seeding density of 5x10^3^ and 5x10^4^ cells/well, respectively), and treated with 200 ng/mL or 500 ng/mL of luciferase-pDNA-LNP, respectively. Following 24 hour LNP treatment, 10% cell counting kit-8 (CCK-8, Enzo Life Sciences) reagent in complete medium was added to cells for 30 minutes incubation and run a colorimetric assay to measure cell viability, according to manufacturer’s protocol. To test the transfection efficiency of luciferase-pDNA-LNP, cells were lysed by cell culture lysis reagent and cell lysates were tested using luminometer upon adding luciferase assay substrate.

MEFs were seeded in ibidi μ-slide 8 well glass bottom chamber (Fitchburg, WI) at the density of 2x10^4^ cells/well. 200 ng/mL of eGFP pDNA-LNP or Lipofectamine were added in complete medium and incubated for 24 hours. eGFP-positive cells were imaged using Leica Stellaris 5 confocal microscopy (Leica Microsystems).

For the primary neuronal culture, E16-E18 embryos were collected for neuronal culture. Single cells were obtained and seede in 8 well glass bottom chamber at 6x10^4^ cells/well.

Neurobasal Plus medium (Gibco, #A3582901) with B27 supplement (Gibco, #A3582801), 1 x GlutaMax and 1% P/S was used for neuronal maintenance. Replace half of the medium every 3 days. On day 3, 5 uM AraC was used for inhibiting proliferating cells. At day 11, neurons were treated with 200ng/mL of eGFP pDNA-LNP or Lipofectamine, and were imaged 24 hours later.

Cytokine measurements were carried out on cell culture medium from RAW macrophages after 4-hour treatment, with IFN-β ELISA kit (Abcam), according to the manufacturer’s instructions.

### Radiolabeling Biodistribution

For biodistribution studies, LNPs or fibrinogen were traced with In-111 or I-125, respectively. LNPs were formulated as described above with 0.1 mol% of 18:0 PE-DTPA using metal-free buffers. Trace metals were removed from the buffers using a Chelex 100 resin, per manufacturer’s instruction. In-111 was mixed with LNPs at a specific activity of 1 μCi of In-111 per 1 μmol of lipid. The mixture was incubated at room temperature for 30 minutes. Fibrinogen was radiolabeled with I-125 using the Iodogen method. Glass tubes coated with 100μg of Iodogen reagent were incubated with fibrinogen (2 mg/mL) and I-125 (115 μCi per μg protein) for 5 minutes on ice. Unincorporated In-111 or I-125 was removed with a Zeba 7kDa desalting spin column (ThermoFisher Scientific). Thin layer chromatography was used to confirm radiolabeling efficiency. All materials were confirmed to have >90% radiochemical purity prior to use.

To trace distribution, fibrinogen was injected 2 minutes prior to LNP intravenous injection. 30 minutes later, blood and organs were harvested, and the radioactivity was quantified with a gamma counter (Wizard2, PerkinElmer). The gamma data and organ weights were used to calculate the tissue biodistribution injected dose per gram.

### DNA uptake and nuclei colocalization

DNA was labeled with SYBR Green by mixing SYBR Green I Nucleic Acid Gel Stain (Invitrogen, Waltham) at 2000X with 1 mg/mL pDNA at room temperature for 30 minutes. Free dye was removed by Zeba desalting spin column. DNA concentration and fluorescent intensity was confirmed by NanoDrop Microvolume Spectrophotometers (Thermo Scientific). SYBR Green labeled pDNAs were then used to either mix with Lipofectamine 2000 to form lipoplexes or to make pDNA-LNPs following previously mentioned LNP synthesis procedure.

HeLa cells were seeded in ibidi μ-slide 8 well glass bottom chamber at a density of 2x10^4^ cells/well and grew overnight. Hoechst 33258 (Cayman Chemical) was added 10 minutes prior to LNP treatment. Cells were treated with 2000 ng/mL of SYBR Green-labeled pDNA-LNPs or pDNA-Lipofectamine 2000 in OPTI-MEM medium. Time-lapse imaging was acquired under 5% CO_2_ at 37°C to trace DNA uptake and nuclei transport. For colocalization analysis, Fiji ImageJ software is used to perform nuclei mask and raw integrated density of SYBR green signal within the nuclei mask were measured and calculated.

### Nanoparticle Tracking Analysis

mRNA and plasmid DNA was labeled using Label IT® Nucleic Acid Labeling Kit (MoBiTec GmbH, Goettingen, Germany) according to the manufacturer’s protocol. Briefly, nucleic acid was mixed with *Label* IT® Reagent at the (w:v) ratio of 5:2, and reacted at 37 °C for 1 hour. Unreacted dye of DNA was purified using G50 Microspin Purification Columns. The labeled mRNA was purified by ethanol precipitation. The final concentration was determined using a NanoDrop Spectrophotometer.

Cy5 labeled mRNA and Cy3 labeled DNA were used to form LNPs to analyze nucleic acid payload in individual nanoparticles. Briefly, LNPs were diluted with deionized water and analyzed using nanoparticle tracking analysis through a Nanosight NS300 (Malvern Panalytical, Westborough, MA). Light scattering mode was used to determine total LNP concentration. Fluorescent mode was used to determine fluorescent nucleic acid loaded LNP concentration. The size of free fluorescent nucleic acid is below the instrument measurement limit, thus was not reported. Percentage of LNP loaded with nucleic acid was calculated as concentration of fluorescent LNP over that of total LNP.

### Quantitative endosomal escape assessments by galectin-9 reporter cell

HeLa cells expressing mCherry-coupled galectin-9 (mCherry-GAL9) were obtained by transfection using plasmids encoding mCherry-galectin 9 (Addgene, Watertown, MA). The galectin-9 reporter HeLa cells were then cultured in ibidi μ-slide 8 well glass bottom chamber at a density of 2x10^4^ cells/well and grew overnight. Cells were dosed with 400 ng/mL of pDNA-LNPs for 24 hours. After fixation with 4% paraformaldehyde (PFA), cells were stained with DAPI and imaged using confocal microscopy. Within individual cell regions of interest, galectin-9 puncta was analyzed via measuring the integrated density of mCherry signals using Fiji ImageJ software.

### p-STING imaging and quantification

Raw macrophages were seeded in ibidi μ-slide 8 well glass bottom chamber at a density of 2x10^5^ cells/well and grew overnight. 1000 ng/mL of pDNA-LNPs were used to treat cells for 4 hours. After treatment, cells were washed with PBS for three times and fixed with 4% paraformaldehyde. For phosphorylated STING staining, cells were permeabilized in 0.1% Triton for 10 minutes, and stained with primary antibody (Phospho-STING (Ser366), 1:200) in blocking buffer containing 10% bovine serum albumin at 4 °C overnight, followed by a secondary antibody (Alexa Fluor 594-conjugated goat-anti rabbit antibody, 1:500) at room temperature for 2 hours. Cell nuclei were labeled using DAPI. Images were acquired by confocal microscopy and mean fluorescent intensity of individual cells were measured by Fiji ImageJ software.

### In Vivo Imaging System

LNPs were injected intravenously into the BALB/c mice. One dose of 10mg/kg baricitinib was injected intravenously 5 minutes prior to injection of C-178 loaded CTS DNA-LNP. At the time of imaging, mice were intraperitoneally injected with 100 uL of 30 mg/mL D-luciferin sodium salt under 3% isoflurane-induced anesthesia, then placed in an IVIS Spectrum machine (PerkinElmer) belly up and imaged for whole body chemiluminescence every 0.2 minutes with automatically determined exposure time for 10-12 images, until the signal reached the peak intensity.

### Statistics

All results are expressed as mean ± SEM unless specified otherwise. Statistical analyses were performed using GraphPad Prism 8 (GraphPad Software) * denotes p<0.05, ** denotes p<0.01, *** denotes p<0.001, **** denotes p<0.0001.

## Acknowledgements

Research reported in this publication was supported by the Institute for RNA Innovation of the Perelman School of Medicine at the University of Pennsylvania (to J.N.), the PhRMA Foundation Postdoctoral Fellowship in Drug Delivery 2023 PFDL 1008128 (to Z.W), Grant R01- HL-157189 (to V.R.M), Grants R01-HL-153510, R01-HL160694, R01-HL164594 (to J.S.B). K.G. acknowledges support from the Johnson Research Foundation and NIH Shared Instrumentation Grant S10-OD018483. The SV-AUC and light scattering experiments were performed at the Johnson Foundation Biophysical and Structural Biology Core Facility (University of Pennsylvania, Philadelphia PA). The SEC-SAXS data were obtained at 16-ID (LIX) at the National Synchrotron Light Source II, a U.S. Department of Energy (DOE) Office of Science User Facility operated for the DOE Office of Science by Brookhaven National Laboratory under Contract No. DE-SC0012704. This work benefited from the use of the SasView application, originally developed under NSF award DMR-0520547. SasView also contains code developed with funding from the European Union’s Horizon 2020 research and innovation programme under the SINE2020 project, grant agreement No 654000. We thank Electron Microscopy Resource Lab (RRID:SCR_022375) for their help with cryo-EM imaging, and Extracellular Vesicle Core for ZetaView analysis.

