## Supplemental Figures and Table for "Multi-stage-mixing to control the supramolecular structure of lipid nanoparticles, thereby creating a core-then-shell arrangement that improves performance by orders of magnitude"

### Slide 1
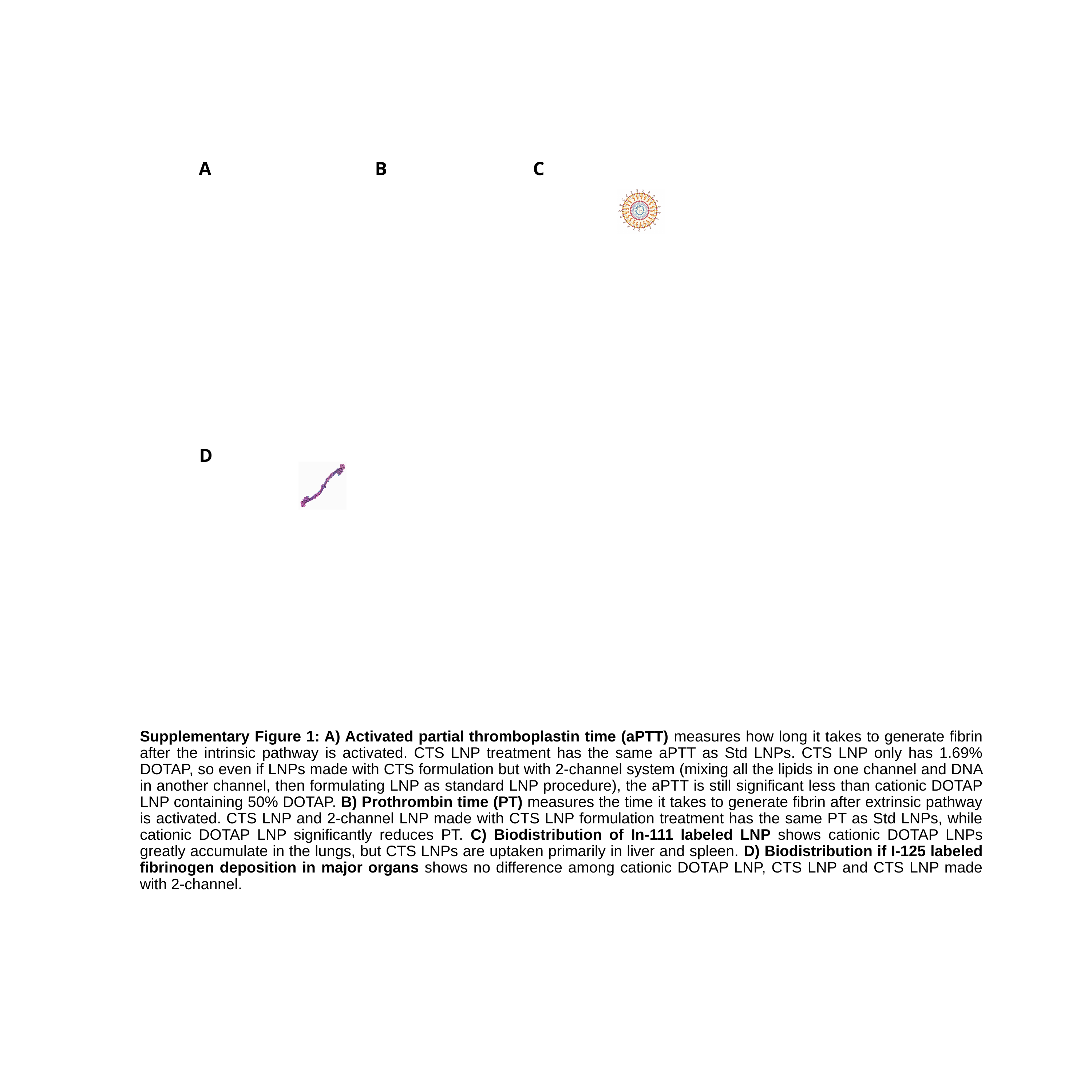

C
A
B
D
Supplementary Figure 1: A) Activated partial thromboplastin time (aPTT) measures how long it takes to generate fibrin after the intrinsic pathway is activated. CTS LNP treatment has the same aPTT as Std LNPs. CTS LNP only has 1.69% DOTAP, so even if LNPs made with CTS formulation but with 2-channel system (mixing all the lipids in one channel and DNA in another channel, then formulating LNP as standard LNP procedure), the aPTT is still significant less than cationic DOTAP LNP containing 50% DOTAP. B) Prothrombin time (PT) measures the time it takes to generate fibrin after extrinsic pathway is activated. CTS LNP and 2-channel LNP made with CTS LNP formulation treatment has the same PT as Std LNPs, while cationic DOTAP LNP significantly reduces PT. C) Biodistribution of In-111 labeled LNP shows cationic DOTAP LNPs greatly accumulate in the lungs, but CTS LNPs are uptaken primarily in liver and spleen. D) Biodistribution if I-125 labeled fibrinogen deposition in major organs shows no difference among cationic DOTAP LNP, CTS LNP and CTS LNP made with 2-channel.

### Slide 2
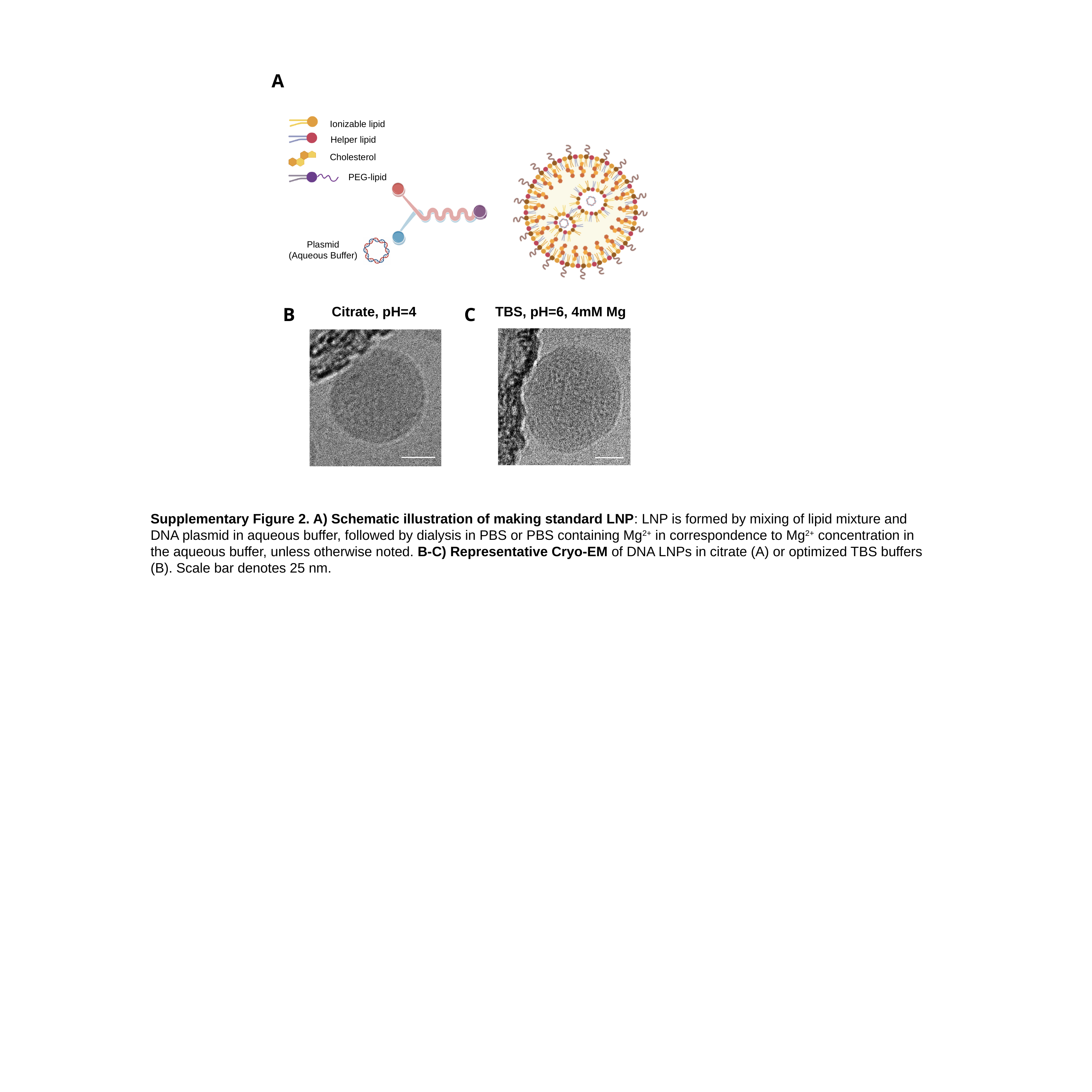

Plasmid
(Aqueous Buffer)
Cholesterol
Helper lipid
Ionizable lipid
PEG-lipid
A
C
B
Citrate, pH=4
TBS, pH=6, 4mM Mg
Supplementary Figure 2. A) Schematic illustration of making standard LNP: LNP is formed by mixing of lipid mixture and DNA plasmid in aqueous buffer, followed by dialysis in PBS or PBS containing Mg^2+^ in correspondence to Mg^2+^ concentration in the aqueous buffer, unless otherwise noted. B-C) Representative Cryo-EM of DNA LNPs in citrate (A) or optimized TBS buffers (B). Scale bar denotes 25 nm.

### Slide 3
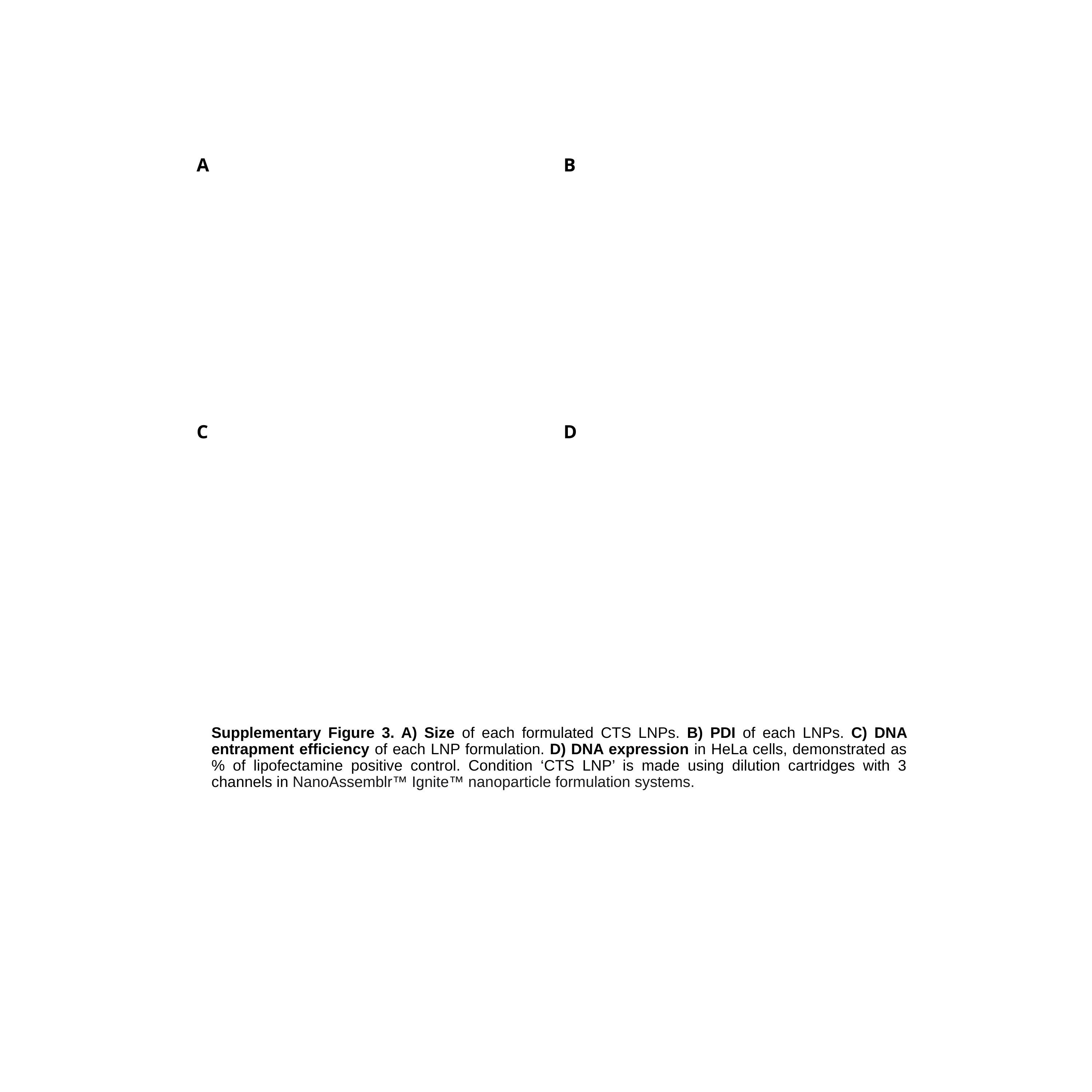

B
A
D
C
Supplementary Figure 3. A) Size of each formulated CTS LNPs. B) PDI of each LNPs. C) DNA entrapment efficiency of each LNP formulation. D) DNA expression in HeLa cells, demonstrated as % of lipofectamine positive control. Condition ‘CTS LNP’ is made using dilution cartridges with 3 channels in NanoAssemblr™ Ignite™ nanoparticle formulation systems.

### Slide 4
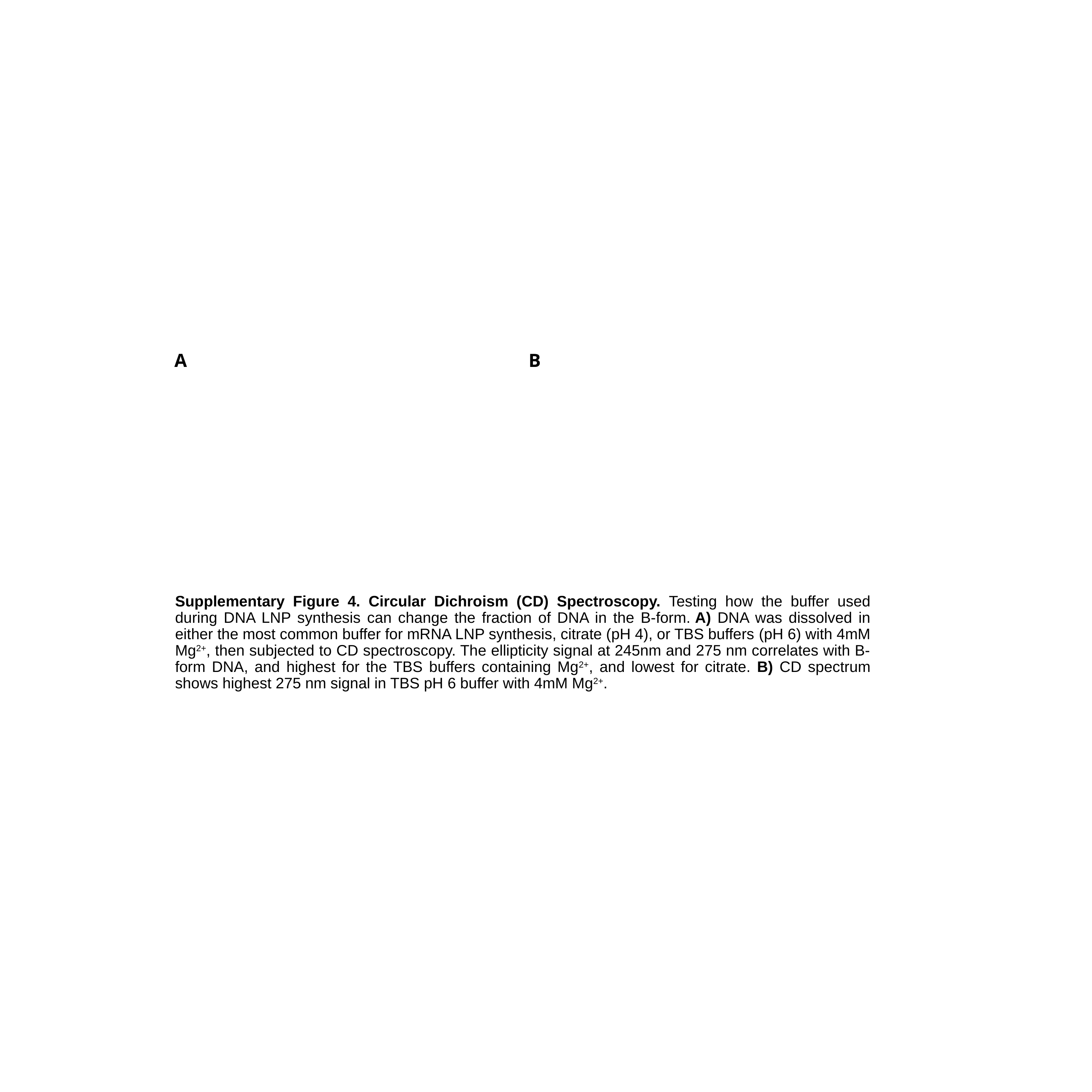

B
A
Supplementary Figure 4. Circular Dichroism (CD) Spectroscopy. Testing how the buffer used during DNA LNP synthesis can change the fraction of DNA in the B-form. A) DNA was dissolved in either the most common buffer for mRNA LNP synthesis, citrate (pH 4), or TBS buffers (pH 6) with 4mM Mg^2+^, then subjected to CD spectroscopy. The ellipticity signal at 245nm and 275 nm correlates with B-form DNA, and highest for the TBS buffers containing Mg^2+^, and lowest for citrate. B) CD spectrum shows highest 275 nm signal in TBS pH 6 buffer with 4mM Mg^2+^.

### Slide 5
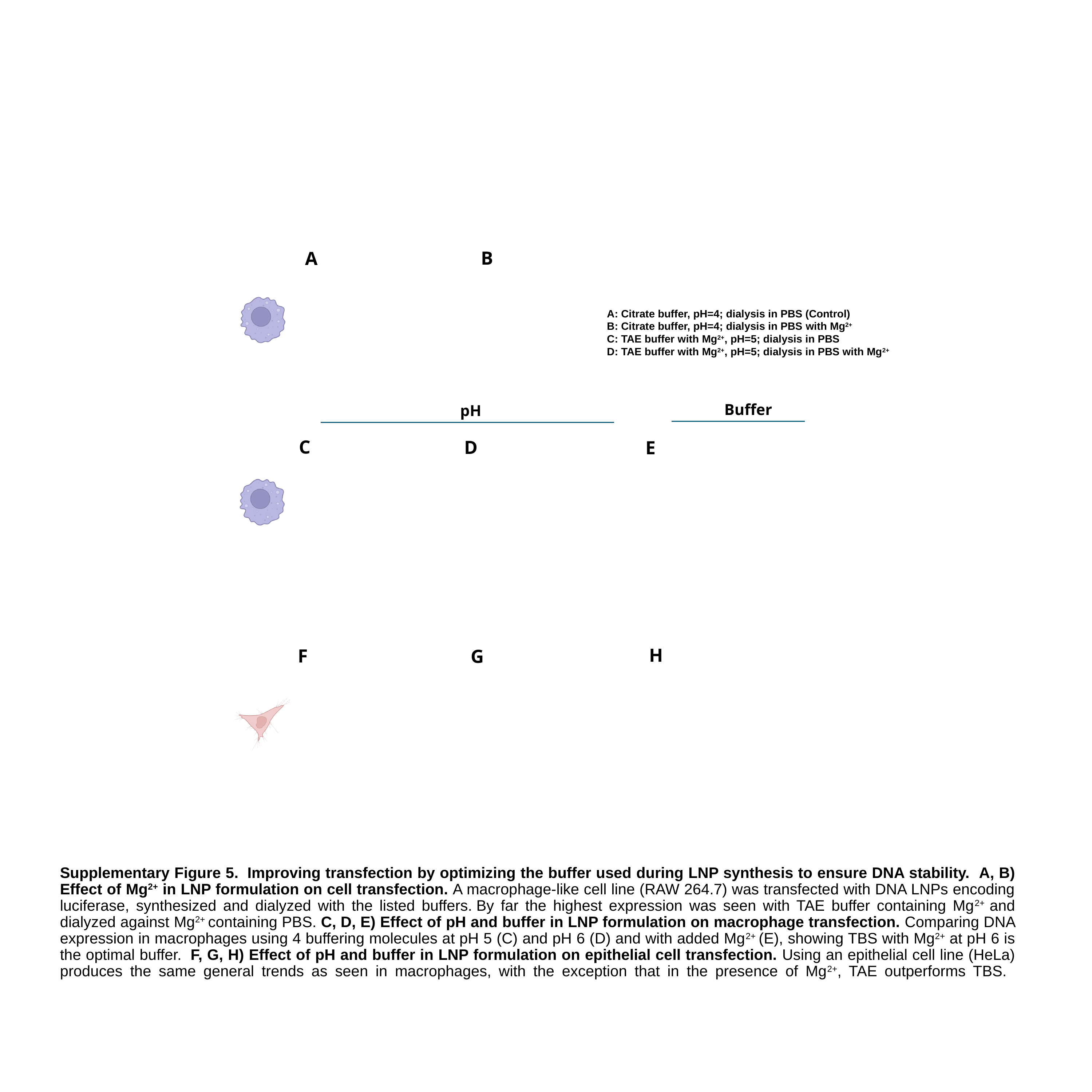

B
A
A: Citrate buffer, pH=4; dialysis in PBS (Control)
B: Citrate buffer, pH=4; dialysis in PBS with Mg^2+^
C: TAE buffer with Mg^2+^, pH=5; dialysis in PBS
D: TAE buffer with Mg^2+^, pH=5; dialysis in PBS with Mg^2+^
Buffer
pH
C
D
E
H
F
G
Supplementary Figure 5.  Improving transfection by optimizing the buffer used during LNP synthesis to ensure DNA stability.  A, B) Effect of Mg^2+^ in LNP formulation on cell transfection. A macrophage-like cell line (RAW 264.7) was transfected with DNA LNPs encoding luciferase, synthesized and dialyzed with the listed buffers. By far the highest expression was seen with TAE buffer containing Mg^2+^ and dialyzed against Mg^2+^ containing PBS. C, D, E) Effect of pH and buffer in LNP formulation on macrophage transfection. Comparing DNA expression in macrophages using 4 buffering molecules at pH 5 (C) and pH 6 (D) and with added Mg^2+^ (E), showing TBS with Mg^2+^ at pH 6 is the optimal buffer.  F, G, H) Effect of pH and buffer in LNP formulation on epithelial cell transfection. Using an epithelial cell line (HeLa) produces the same general trends as seen in macrophages, with the exception that in the presence of Mg^2+^, TAE outperforms TBS.

### Slide 6
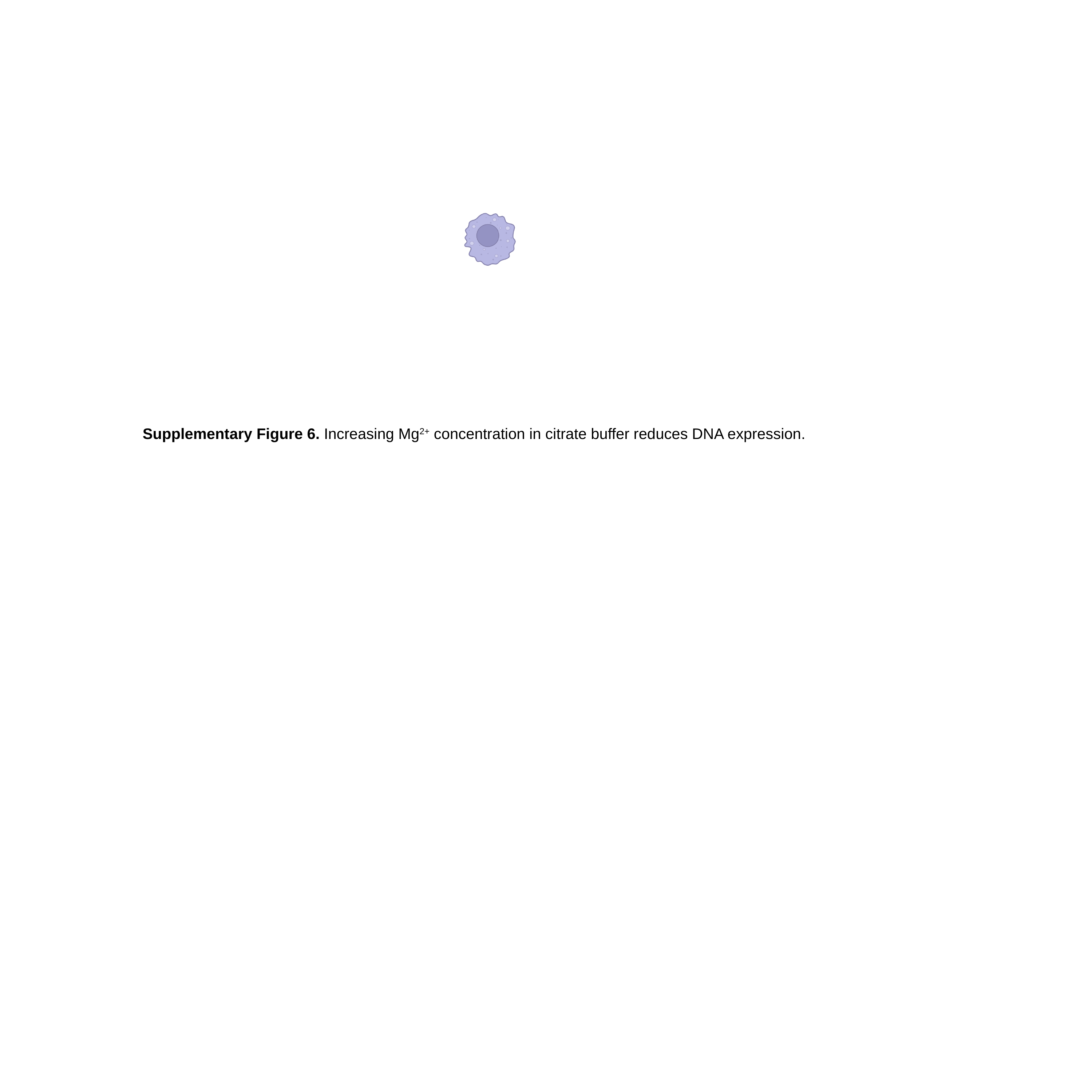

Supplementary Figure 6. Increasing Mg^2+^ concentration in citrate buffer reduces DNA expression.

### Slide 7
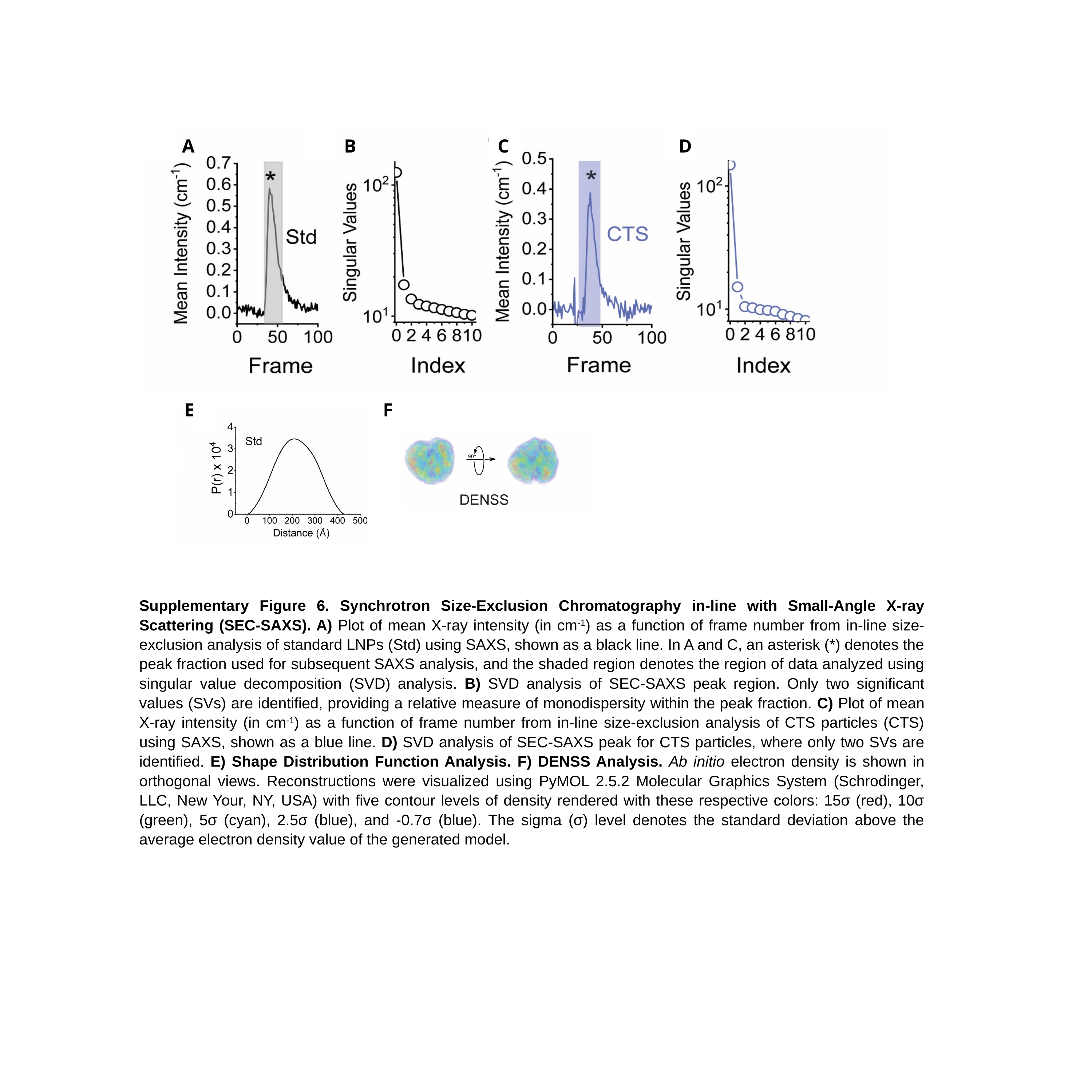

A
B
D
C
F
E
Supplementary Figure 6. Synchrotron Size-Exclusion Chromatography in-line with Small-Angle X-ray Scattering (SEC-SAXS). A) Plot of mean X-ray intensity (in cm^-1^) as a function of frame number from in-line size-exclusion analysis of standard LNPs (Std) using SAXS, shown as a black line. In A and C, an asterisk (*) denotes the peak fraction used for subsequent SAXS analysis, and the shaded region denotes the region of data analyzed using singular value decomposition (SVD) analysis. B) SVD analysis of SEC-SAXS peak region. Only two significant values (SVs) are identified, providing a relative measure of monodispersity within the peak fraction. C) Plot of mean X-ray intensity (in cm^-1^) as a function of frame number from in-line size-exclusion analysis of CTS particles (CTS) using SAXS, shown as a blue line. D) SVD analysis of SEC-SAXS peak for CTS particles, where only two SVs are identified. E) Shape Distribution Function Analysis. F) DENSS Analysis. Ab initio electron density is shown in orthogonal views. Reconstructions were visualized using PyMOL 2.5.2 Molecular Graphics System (Schrodinger, LLC, New Your, NY, USA) with five contour levels of density rendered with these respective colors: 15σ (red), 10σ (green), 5σ (cyan), 2.5σ (blue), and -0.7σ (blue). The sigma (σ) level denotes the standard deviation above the average electron density value of the generated model.

### Slide 8
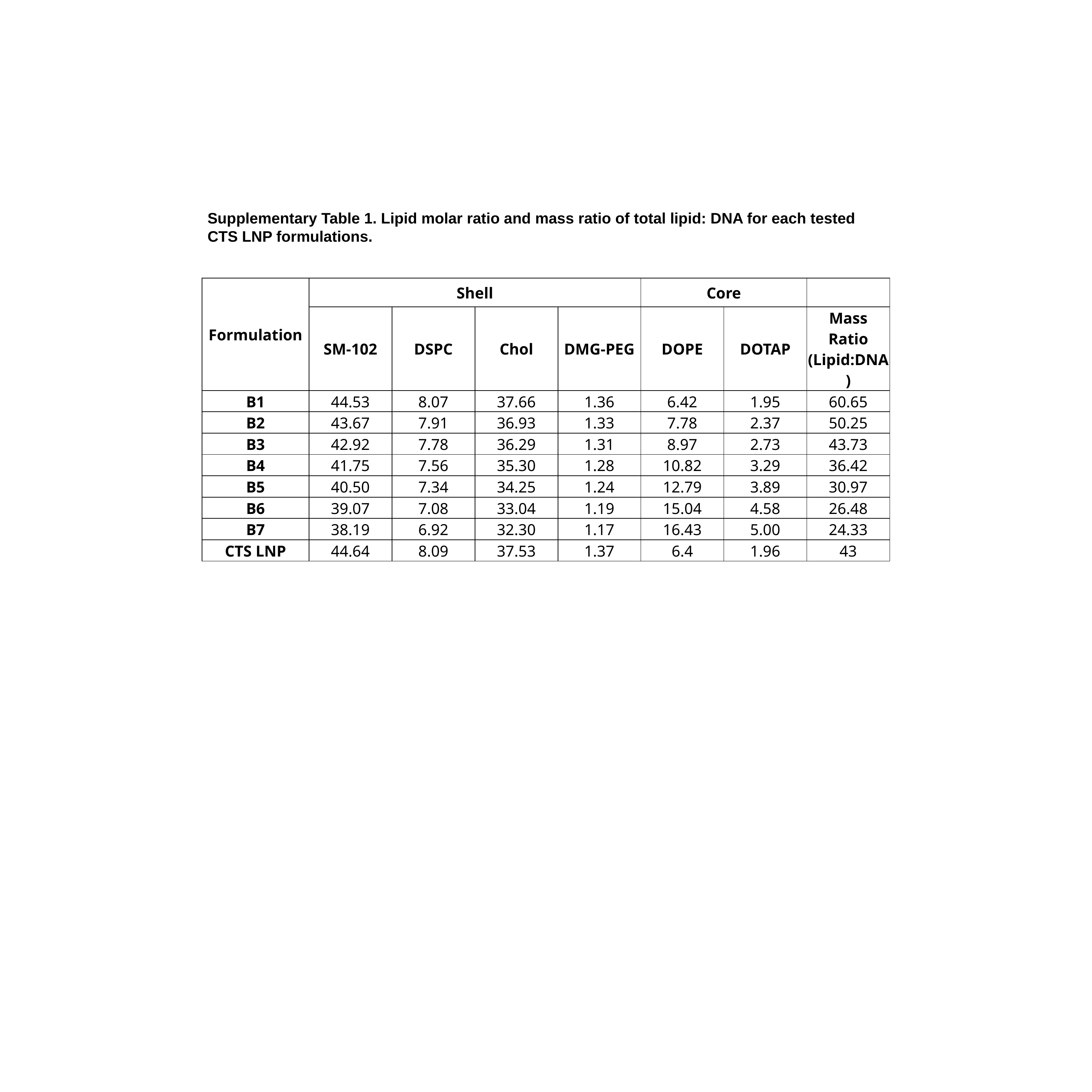

Supplementary Table 1. Lipid molar ratio and mass ratio of total lipid: DNA for each tested CTS LNP formulations.
| Formulation | Shell | | | | Core | | |
| --- | --- | --- | --- | --- | --- | --- | --- |
| | SM-102 | DSPC | Chol | DMG-PEG | DOPE | DOTAP | Mass Ratio (Lipid:DNA) |
| B1 | 44.53 | 8.07 | 37.66 | 1.36 | 6.42 | 1.95 | 60.65 |
| B2 | 43.67 | 7.91 | 36.93 | 1.33 | 7.78 | 2.37 | 50.25 |
| B3 | 42.92 | 7.78 | 36.29 | 1.31 | 8.97 | 2.73 | 43.73 |
| B4 | 41.75 | 7.56 | 35.30 | 1.28 | 10.82 | 3.29 | 36.42 |
| B5 | 40.50 | 7.34 | 34.25 | 1.24 | 12.79 | 3.89 | 30.97 |
| B6 | 39.07 | 7.08 | 33.04 | 1.19 | 15.04 | 4.58 | 26.48 |
| B7 | 38.19 | 6.92 | 32.30 | 1.17 | 16.43 | 5.00 | 24.33 |
| CTS LNP | 44.64 | 8.09 | 37.53 | 1.37 | 6.4 | 1.96 | 43 |
